# Chronic stress induces sex-specific functional and morphological alterations in cortico-accumbal and cortico-tegmental pathways

**DOI:** 10.1101/2020.09.22.306860

**Authors:** Thibault P. Bittar, Mari Carmen Pelaez, Jose Cesar Hernandez Silva, Francis Quessy, Andrée-Anne Lavigne, Daphnée Morency, Léa-Jeanne Blanchette, Eric Arsenault, Yoan Cherasse, Josée Seigneur, Igor Timofeev, Chantelle F. Sephton, Christophe D. Proulx, Benoit Labonté

## Abstract

**Background:** The medial prefrontal cortex (mPFC) is part of a complex circuit controlling stress responses by sending projections to different limbic structures including the nucleus accumbens (NAc) and ventral tegmental area (VTA). However, the impact of chronic stress on NAc- and VTA-projecting mPFC neurons is still unknown and the distinct contribution of these pathways to stress responses in males and females is unclear.

**Methods:** Behavioral stress responses were induced by 21 days of chronic variable stress (CVS) in male and female C57BL6 mice. An inter-sectional viral approach was used to label both pathways and assess the functional, morphological, and transcriptional adaptations in NAc- and VTA-projecting mPFC neurons in stressed males and females. Using chemogenetic approaches, we modified neuronal activity of NAc-projecting mPFC neurons to decipher their contribution to stress phenotypes.

**Results:** CVS induced depressive-like behaviors in males and females. NAc- and VTA-projecting mPFC neurons exhibited sex-specific functional, morphological, and transcriptional alterations. The functional changes were more severe in females in NAc-projecting mPFC neurons while males exhibited more drastic reductions in dendritic complexity in VTA-projecting mPFC neurons after CVS. Finally, chemogenetic overactivation of the cortico-accumbal pathway triggered anxiety and behavioral despair in both sexes while its inhibition rescued the phenotype only in females.

**Conclusions:** Our results suggest that by changing the activity of transcriptional programs controlling neuronal plasticity, CVS interferes with the morphological and synaptic properties of the cortico-accumbal and tegmental pathways differently in males and females contributing to the expression of anxiety and depressive-like behaviors distinctly in a sex-specific fashion.

## Introduction

Major depressive disorder (MDD) is a debilitating psychiatric disorder affecting more than 300 million people worldwide (1). According to the World Health Organization, MDD is the leading cause of disability, imposing major economic, medical, and human burdens on societies (2). Amongst the different brain regions affected in MDD patients, the medial prefrontal cortex (mPFC) emerges as a hub region (3). Imaging studies on treatment-resistant MDD patients revealed elevated metabolic activity in the mPFC (4,5). Decreased dendritic complexity and spine density in the mPFC has also been reported in MDD post-mortem tissue (6,7) and mouse models of depressive-like behaviors (8–10). Furthermore, recent transcriptional analyses highlighted a major reorganization of transcriptional structures in the mPFC of men and women with MDD with different mouse models of chronic stress reproducing a significant proportion of these changes (11–13). While consolidating the morpho-functional and molecular impact of chronic stress on the mPFC, these findings do not address how the different neuronal circuits controlling the expression of stress responses in males and females are affected.

Substantial lines of evidence suggest that cortically-driven dysregulation of subcortical circuits underlies the expression of depressive behaviors in humans and stress-induced depressive-like behaviors in animals (14–18). The mPFC projects densely to the nucleus accumbens (NAc) and the ventral tegmental area (VTA), two brain regions that have been consistently associated with the expression of MDD in humans and stress responses in mouse models of depression (19– 23). The NAc is involved in reward processing by action selection and motivated behavioral response (24,25) while the VTA is playing a major role in the processing of appetitive and aversive stimuli as part of the reward circuit (26,27). Moreover, functional studies showed that the optogenetic activation of the cortico-accumbal pathway reverses social avoidance induced by chronic social defeat (20) while acute stimulations of mPFC nerve terminals in the NAc rescue cholecystokinin-induced social avoidance and anhedonia in male mice (28). Elevated activity of the cortico-accumbal pathway has also been described during pro-hedonic reward-seeking in mice (29). Conversely, other studies showed that continuous stimulation of the mPFC nerve terminal in the NAc decreases social preference (14) and hyperexcitability of the NAc-projecting mPFC neurons was shown to suppress reward-motivated behaviors in male rats (15). In contrast, while a few studies highlighted the contribution of the projections from VTA to mPFC in modulating susceptibility to stress (23,30,31), the role of the cortico-tegmental pathway in controlling stress responses in males and females is still poorly understood.

Here, we used a multi-dimensional approach to define the sex-specific morpho-functional impact of chronic variable stress on cortico-accumbal and cortico-tegmental pathways and elaborated on the specific contribution of the cortico-accumbal pathway in mediating depressive-like behaviors in males and females.

## Methods and materials

**Detailed Methods** are provided in **Supplemental Information**. All animal experiments were performed in male and female C57BL/6NCrl mice aged 8 to 14 weeks old. Depressive-like behaviors were induced through chronic variable stress (CVS) as previously described (11,32).

For electrophysiological recordings, mice were injected with AAV2-retro-EF1a-FlpO in the NAc, AAV2-retro-hSyn-Cre in the VTA, and AAV-EF1a-fDIO-mCherry/AAV-hSyn-DIO-EGFP in the mPFC. After 21 days of CVS, whole-cell voltage-clamp recordings were performed to collect passive membrane properties and spontaneous excitatory and inhibitory postsynaptic currents (respectively sEPSCs and sIPSCs) in layer II/III for NAc-projecting mPFC neurons and layer V for VTA-projecting mPFC neurons in the prelimbic cortex.

For morphological analyses, mice were injected with AAV2-retro-hSyn-Cre in NAc or VTA and AAV-hSyn-DIO-EGFP in the mPFC. After 21 days of CVS, brain tissue was prepared for morphological 3D reconstruction of layer II/III for NAc-projecting mPFC neurons and layer V for VTA-projecting mPFC neurons in the prelimbic cortex and spine detection using Neurolucida 360. Sholl analysis was performed as previously described (33). Automatic spine detection with manual correction was used to identify and quantify spines along the full length of the apical dendritic tree.

Cell type-specific transcriptional assessment was performed on mice injected with AAV2-retro-EF1a-FlpO in the NAc, AAV2-retro-hSyn-Cre in the VTA, and AAV-EF1a-fDIO-mCherry/AAV-hSyn-DIO-EGFP in the mPFC. After 21 days of CVS, mice were sacrificed to collect mPFC tissue for fluorescence-assisted cell sorting (FACS) as previously described (34). RT-qPCR was performed with a QuantStudio 5 Real-Time PCR System. Relative gene expression assessment was performed with GAPDH and 18S as reference genes.

Chemogenetic manipulations were performed as described before (35,36) with the Designer Receptor Exclusively Activated by Designer Drug (DREADD)’s Agonist 21 (Compound 21 (C21)) (37,38). For both acute and sub-chronic chemogenetic activation experiments, mice were injected with AAV-hSyn-DIO-hM3Dq-mCherry in the mPFC and AAV2-retro-CAG-Cre-EGFP-WPRE in the NAc. For the acute experiments, mice were injected intraperitoneally 30 min prior to behavioral testing. For the sub-chronic chemogenetic activation, mice underwent CVS for 3 days combined with intraperitoneal injections twice a day. For the chemogenetic inhibition experiments, mice were injected with AAV-hSyn-DIO-hM4Di-mCherry in the mPFC and AAV2-retro-CAG-Cre-EGFP-WPRE in the NAc before going through 21 days of CVS. 30 min prior to each behavioral assessment, mice were injected intraperitoneal.

Data were analyzed through independent sample t-tests, one-way or two-way ANOVA with Tukey posthoc tests when justified. All statistical analyses were achieved using GraphPad Prism 8.

## Results

In this study, we defined the morphological, functional, and molecular impact of chronic stress on the activity of the NAc- and VTA-projecting mPFC neurons in males and females (**Figure 1**). To do so, we used a model of chronic variable stress (CVS) (32). Our results show that 21 days of CVS induce a complex depressive-like phenotype in both males and females (**Figures 2A-G**). CVS increased the latency to eat in the novelty-suppressed feeding test (NSF; F_(1,28)_=34.470, *p*<0.0001, **Figure 2B**), decreased the time spent in the open arms of the elevated plus-maze test (EPM; F_(1,28)_=55.572, *p*<0.0001, **Figure 2C**), decreased the time spent grooming in the splash test (F_(1,28)_=30.895, *p*<0.0001, **Figure 2D**), decreased the latency to immobility in the forced swim test (FST; F_(1,28)_=44.918, *p*<0.0001, **Figure 2E**) and decreased preference for sweet water in the sucrose preference test (F_(1,28)_=26,745, *p*<0.0001, **Figure 2F**). No sex-specific effect was observed in any of the tests. We further confirmed the consistency of our behavioral assessment showing that CVS increases overall behavioral emotionality (39) in both sexes (F_(1,28)_=102.95, *p*<0.0001, **Figure 2G**).

**Figure 1.**
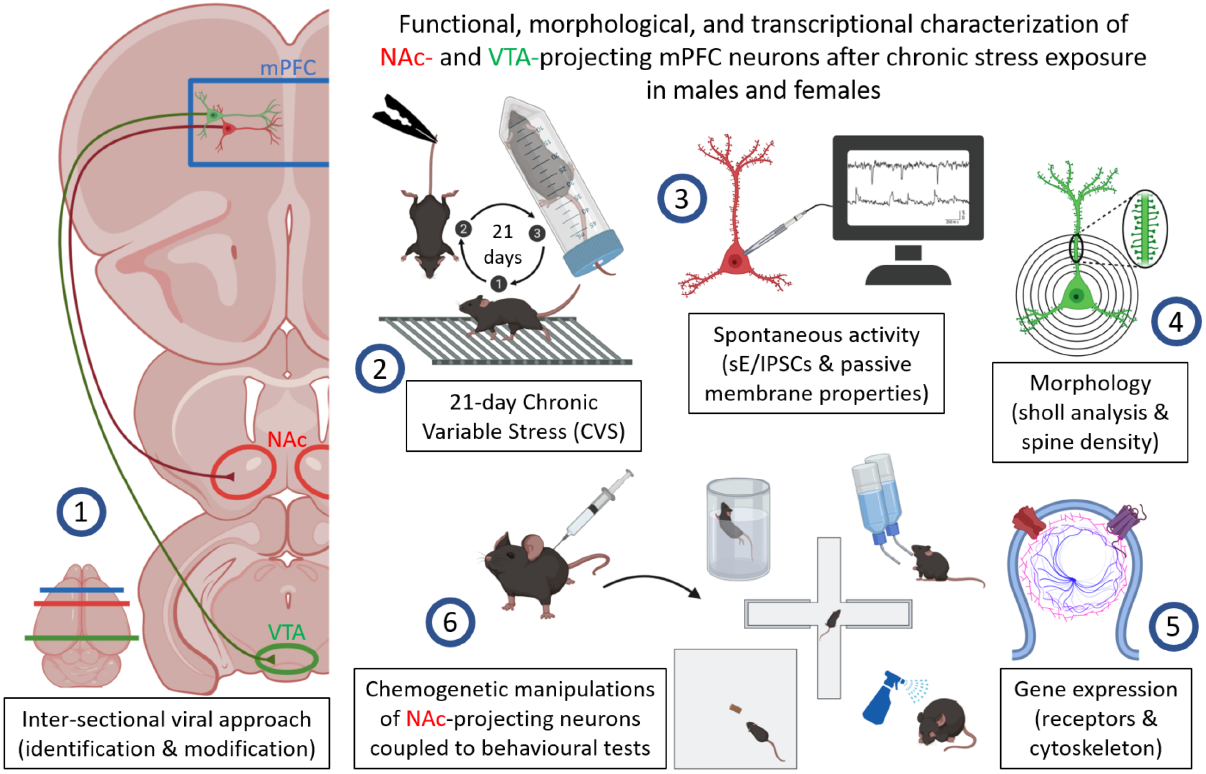
Multiscale analysis of functional, morphological, and transcriptional alterations of the cortico-accumbal and cortico-tegmental pathways after chronic stress exposure in males and females. (1) Inter-sectional viral approaches were used to label NAc- and VTA-projecting mPFC neurons and then express a transgene. (2) Mice underwent a 21-day chronic variable stress (CVS) protocol to induce depressive-like behaviors. (3) Spontaneous excitatory and inhibitory postsynaptic currents and passive membrane properties were recorded. (4) Neuronal morphology and spine density were assessed. (5) Differential expression of genes coding for receptors and proteins involved in cytoarchitecture and neuronal plasticity was assessed. (6) Chemogenetic manipulations of NAc-projecting mPFC neurons were used to determine the behavioral contribution of this pathway in both sexes.

**Figure 2.**
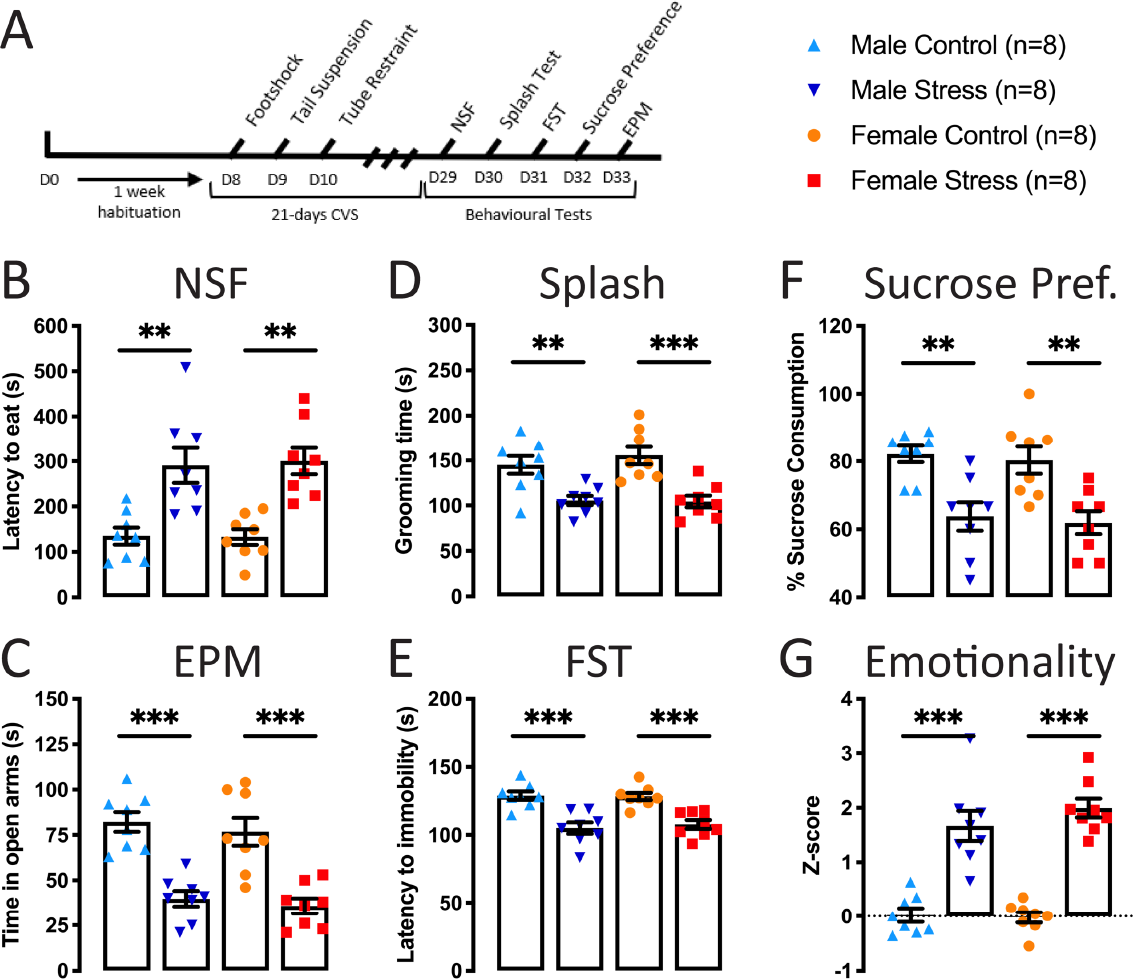
CVS induces anxiety, behavioral despair, and anhedonia in male and female mice. **(A)** Schematic representation of the CVS protocol followed by the behavioral tests used to assess depressive-like behaviors in males and females. **(B-C)** Anxiety was assessed with the novelty-suppressed feeding test (NSF) and the elevated plus-maze test (EPM). **(B)** The latency to eat in the NSF is increased after CVS and **(C)** the time spent in the open arms of the EPM is reduced in both sexes. **(D-E)** Behavioral despair was assessed with the splash test and the forced swim test (FST). **(D)** The time spent grooming and **(E)** the latency to immobility in the FST is decreased after CVS in both sexes. **(F)** Anhedonia was assessed with the sucrose preference test. Sucrose preference was significantly decreased after CVS in both sexes. **(G)** Overall emotionality is increased in both sexes after 21-day CVS. Significance was determined using two-way ANOVA with posthoc Tukey multiple comparisons tests. Symbols and bars represent the group average ± SEM; ** *p* < 0.01, *** *p* < 0.005.

### Mapping the mPFC neuronal pathways projecting to the NAc and VTA

In order to evaluate the impact of CVS on NAc- and VTA-projecting mPFC neuronal pathways, we first tested our capacity to accurately label and target these two pathways in the mouse brain. We used an inter-sectional viral approach to express red and green fluorescent reporters in the mPFC neurons projecting, respectively, to the NAc and the VTA (**Figure 3A**). Our results show that NAc- and VTA-projecting mPFC neurons follow a specific topological organization with a large proportion of the neurons located in layer II/III projecting to NAc (81.3%) while the neurons in layer V are mainly VTA-projecting neurons (73.1%; **Figure 3B**). Additionally, consistent with a previous report (40), we found some double-labeled neurons in layers II/III and V suggesting that a small proportion of cells from the mPFC (7.3%) are sending collaterals to the NAc and the VTA.

**Figure 3.**
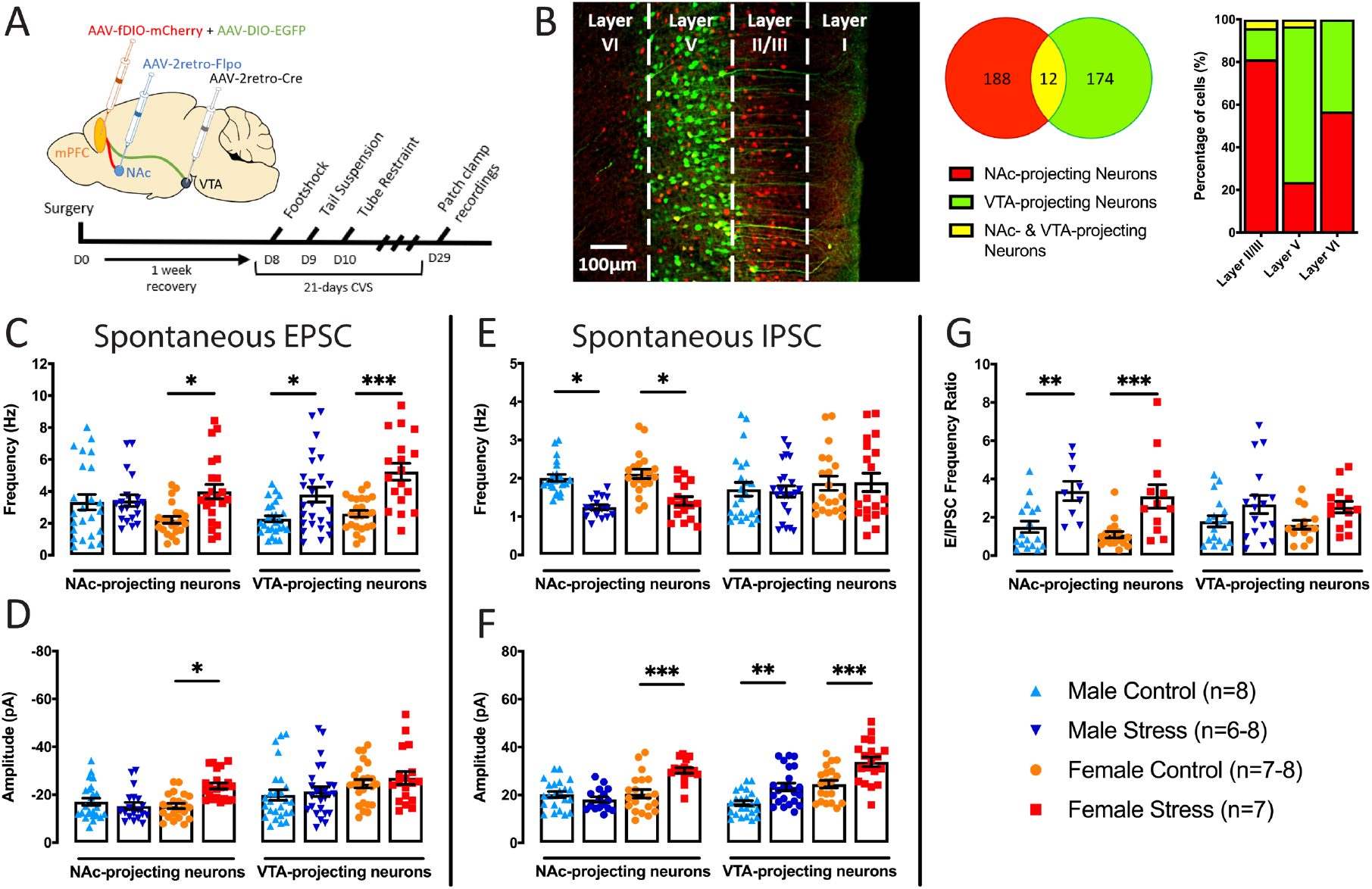
CVS alters NAc- and VTA-projecting mPFC neuron functional activity in a sex-specific fashion. **(A)** Schematic representation of the viral injections and the experimental timeline. **(B)** Representative viral infection showing NAc- and VTA-projecting mPFC. The NAc-projecting neurons are labeled in red and are mainly localized on layer II/III. The VTA-projecting neurons are labeled in green and are mainly localized on the layer V. The Venn diagram represents the count of both neuronal populations and the bar plot shows the cortical layer representation of both neuronal populations. **(C)** Frequency of spontaneous EPSC in NAc- and VTA-projecting mPFC neurons in male (blue) and females (red). **(D)** Amplitude of spontaneous EPSC in NAc- and VTA-projecting neurons in male (blue) and females (red). **(E)** Frequency of spontaneous IPSCs in NAc- and VTA-projecting mPFC neurons in males (blue) and females (red). **(F)** Amplitude of spontaneous IPSCs in NAc- and VTA-projecting mPFC neurons in males (blue) and females (red). **(G)** E/IPSCs frequency ratio in NAc- and VTA-projecting mPFC neurons in males (blue) and females (red). Significance was determined using two-way ANOVA with posthoc Tukey multiple comparisons tests. Symbols and bars represent the group average ± SEM; * *p* < 0.05, ** *p* < 0.01, *** *p* < 0.005.

### CVS induces sex-specific functional changes in NAc- and VTA-projecting mPFC neuronal pathways

To gain insights into the functional impact of CVS on NAc- and VTA-projecting mPFC neuronal pathways, we measured variations in the electrophysiological properties of mPFC glutamatergic pyramidal neurons projecting to either pathway. We first labeled both neuronal populations using the same trans-sectional viral approach described above (**Figure 3A**) before stressing mice through 21 days of CVS and recording spontaneous activity for excitatory and inhibitory postsynaptic currents in males and females (**Supplemental Figures S1A-B**).

Our analysis shows that CVS increases the frequency of sEPSCs in NAc- and VTA-projecting mPFC neurons in females (NAc-projecting: F_(1,83)_=5.3706, *p*=0.0229, VTA-projecting: F_(1,90)_=32.417, *p*<0.0001) but only in VTA-projecting mPFC neurons in males (**Figure 3C**). Our results also show that CVS, in females but not males, increases the amplitude of sEPSCs in NAc-projecting mPFC neurons (F_(1,83)_=6.1401, *p*=0.0152) with no change in VTA projections (F_(1,90)_=0.78956, *p*=0.3766, **Figure 3D**). In contrast, our findings suggest that CVS decreases the frequency of sIPSCs in NAc-projecting mPFC neurons from both males and females (F_(1,71)_=50.087, *p*<0.0001) with no effect in VTA-projecting mPFC neurons (**Figure 3E**). Our analysis also revealed that CVS increases the amplitude of sIPSCs in NAc- and VTA-projecting mPFC neurons in females (NAc: F_(1,75)_=7.3147, p=0.0085 and VTA: F_(1,81)_=26.553, *p*<0.0001) but only in VTA-projecting mPFC neurons in males (VTA: *p*=0.0102; **Figure 3F**). Interestingly, analysis of the E/IPSC frequency ratio suggests that CVS shifts the excitatory/inhibitory balance toward excitation in the cortico-accumbal pathway of males and females but not in the cortico-tegmental pathway (NAc: F_(1,52)_=23.34, *p*<0.0001; VTA: F_(1,52)_=1.9581, *p*=0.1654; **Figure 3G**). Finally, our analyses of NAc-projecting mPFC neurons revealed a significant decrease in membrane resistance in stressed females but not in males (F_(1,83)_=10.176, *p*=0.002) although no changes were observed in membrane capacitance in both sexes (**Supplemental Figures S1C-D**). Also, CVS has no impact on passive membrane resistance and capacitance in males or females in VTA-projecting mPFC neurons.

Together, our results suggest that 21 days of CVS changes the excitatory and inhibitory balance in NAc- and VTA-projecting mPFC neurons by altering excitatory and inhibitory inputs to these neuronal populations in a sex-specific fashion.

### Sex-specific morphological adaptations to CVS in NAc- and VTA-projecting mPFC neurons

Next, we tested whether CVS induces pathway-specific morphological adaptations in males and females. To do so, we used our inter-sectional viral approach to express EGFP in mPFC neurons projecting either to the NAc or the VTA (**Figure 4A**). Following 21 days of CVS, brain samples were processed for confocal imaging and 3D neuronal reconstruction of NAc- and VTA-projecting glutamatergic pyramidal mPFC neurons.

**Figure 4.**
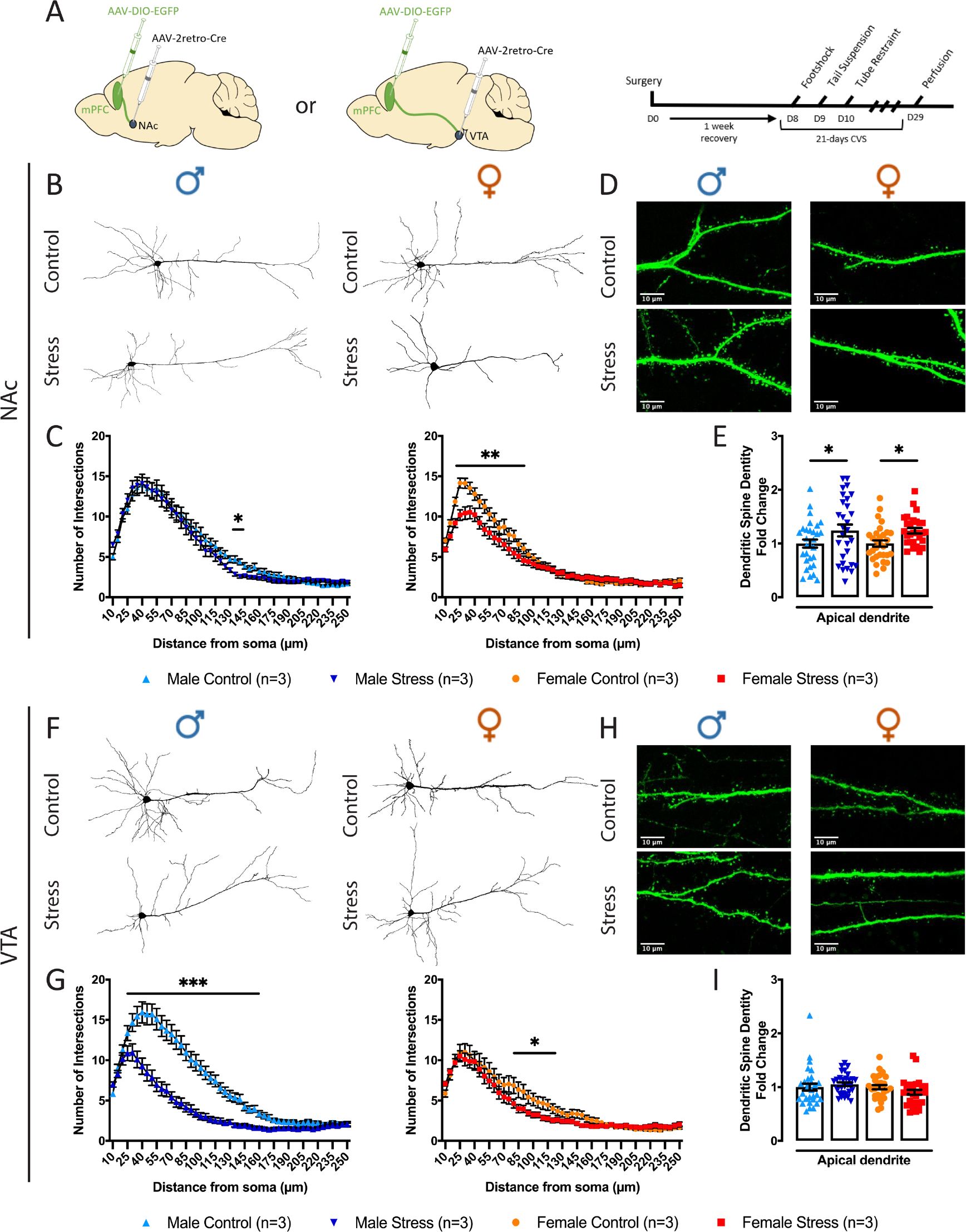
NAc- and VTA-projecting mPFC neurons morphology is altered differently in males and females after chronic stress exposure. **(A)** Schematic representation of the viral injections and experimental timeline. **(B)** Representative examples of 3D-reconstructed neurons for control and stressed males and females in NAc-projecting mPFC neurons. **(C)** Sholl analyses of the number of intersections relative to the distance from the soma for both apical and basal dendritic branches of NAc-projecting mPFC neurons in males and females. **(D-E)** Total spine densities in NAc-projecting mPFC neurons in males (blue) and females (red) and representative examples. **(F)** Representative examples of 3D-reconstructed neurons for control and stressed males and females in VTA-projecting mPFC neurons. **(G)** Sholl analyses of the number of intersections relative to the distance from the soma for both apical and basal dendritic branches of VTA-projecting mPFC neurons in males and females. **(H-I)** Total spine densities in VTA-projecting mPFC neurons in males (blue) and females (red) and representative examples. Significance was determined using mixed-effects models for repeated measures with posthoc Sidak multiple comparisons test for Sholl analyses and two-way ANOVA with posthoc Tukey multiple comparisons test for spine density. Symbols and bars represent the group average ± SEM; * *p* < 0.05, ** *p* < 0.01, *** *p* < 0.005.

Our analyses of NAc-projecting mPFC neurons revealed a significant decrease in the total number of intersections in the medial section in relation to the soma of stressed male dendritic trees (Total: F_(48,2744)_=1.3794, *p*=0.0435; Apical: F_(48,2744)_=1.8402, *p*=0.0004; Basal: F_(30,1263)_=1.8754, *p*=0.0029; **Figures 4B-C** and **Supplemental Figures S2A-B**) and in the proximal section for stressed females (Total: F_(48,2437)_=3.431, *p*<0.0001; Apical: F_(48,2437)_=2.0798, *p*<0.0001; Basal: F_(30,1264)_=1.500, *p*=0.0410). At the spine level, our analysis of NAc-projecting mPFC neurons revealed an increase in the total spine density along the full apical dendrite of both stressed males and females (F_(1,113)_=9.5569, *p*=0.0025, **Figures 4D-E**).

Striking differences were observed in the VTA-projecting mPFC neurons. Indeed, our analyses revealed a drastic decrease in total number of intersections in stressed males (F_(48,2713)_=15.25, *p*<0.0001), consistent across apical (F_(48,2709)_=2.9527, *p*<0.0001) and basal (F_(30,1223)_=9.1279, *p*<0.0001) sections (**Figures 4F-G** and **Supplemental Figures S2C-D)**. In contrast, CVS in females induced a much subtler decrease in the number of intersections (F_(48,2716)_=1.889, *p*=0.0002) restricted to basal dendritic trees (F_(30,1124)_=1.785, *p*=0.006) and not in the apical (F_(48,2710)_=1.1888, *p*=0.1765). These changes in the dendritic complexity of VTA-projecting mPFC neurons induced by CVS in males and females were, however, not associated with major changes in spine density along the full apical dendrite length (F_(1,115)_=0.19154, *p*=0.6625; **Figures 4H-I**).

Overall, these results suggest that CVS induces a consistent reorganization of mPFC neurons morphological properties, with VTA-projecting mPFC neurons more affected in males, and NAc-projecting more affected in females.

### CVS induces pathway-specific transcriptional changes in males and females

Our analyses showed that CVS induces functional and morphological changes in NAc- and VTA-projecting mPFC neurons in a sex-specific fashion. Next, we wanted to test whether these functional and morphological adaptations associate with transcriptional changes. To do so, we identified a panel of genes controlling various aspects of cytoskeletal plasticity and neuronal activity (41,42) previously reported to be differentially expressed in the mPFC following chronic stress in mice (43–46) and MDD in human post-mortem tissue (47–50).

We first labelled both neuronal populations using our inter-sectional viral approach to express mCherry and EGFP in NAc- and VTA-projecting mPFC neurons, respectively (**Figure 5A**). Mice were then stressed for 21 days and both neuronal populations were isolated through FACS with a high level of purity (**Figure 5B)**. In NAc-projecting mPFC neurons, our analyses shows that CVS increases significantly *Grin1* (F_(1,20)_=9.395, *p*=0.0061), *Grin2a* (F_(1,20)_=6.914, *p*=0.0161), *Gabra2* (F_(1,20)_=5.091, *p*=0.0354), *Shank1* (F_(1,20)_=14.91, *p*=0.001), and *Dgl4* (F_(1,20)_=14.11, *p*=0.0012) and decreases *Homer1* (F_(1,20)_=9.185, *p*=0.0066) and *Actb* (F_(1,20)_=13.57, *p*=0.0015) expressions in females but not males (**Figure 5C**). In contrast, CVS significantly increases *Actb* (F_(1,20)_=13.57, *p*=0.0015) and decreases *Shank1* (F_(1,20)_=14.91, *p*=0.001) expression in males (**Figure 5C**). The effect of CVS in VTA projecting mPFC neurons were far more diverse. For instance, our analyses show that CVS increases *Grin2a* (F_(1,20)_=8.533, *p*=0.0084), *Gabra1* (F_(1,20)_=23.58, *p*<0.0001), *Gabra2* (F_(1,20)_=20.86, *p*=0.0002), *Homer1* (F_(1,20)_=17.34, *p*=0.0005), *Shank1* (F_(1,20)_=15.75, *p*=0.0008) and *Dlg4* (F_(1,20)_=20.77, *p*=0.0002) and decreases *Grin1* expression (F_(1,20)_=16.81, *p*=0.0006) in females (**Figure 5D**) while in males, CVS increases *Gabra1* (F_(1,20)_=23.58, *p*<0.0001), *Homer1* (F_(1,20)_=17.34, *p*=0.0005) and *Shank1* (F_(1,20)_=15.75, *p*=0.0008) and decreases *Dlg4* (F_(1,20)_=20.77, *p*=0.0002) and *Actb* (F_(1,20)_=11.33, *p*=0.0031) gene expression (**Figure 5D**).

**Figure 5.**
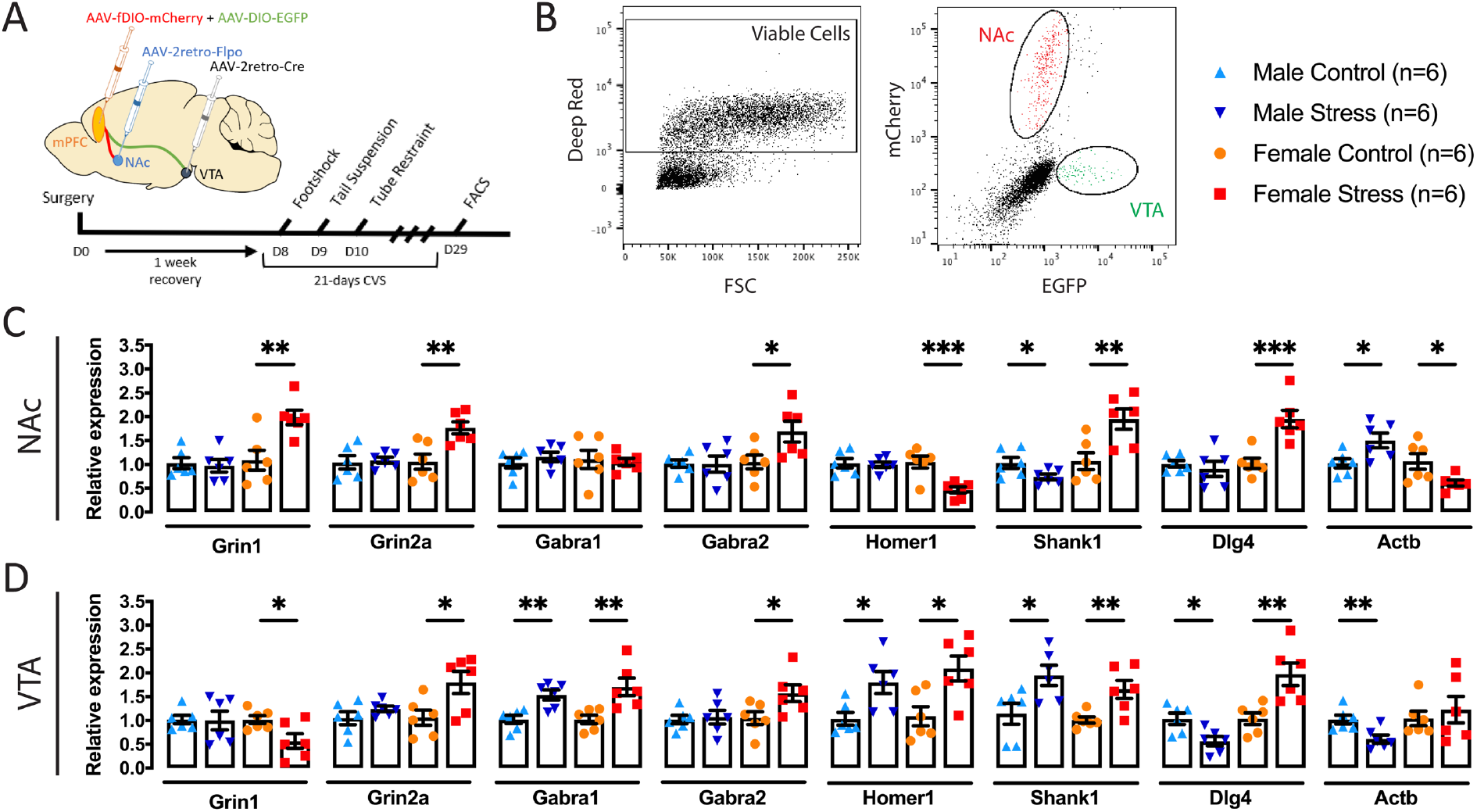
CVS induces pathway-specific transcriptional changes in males and females. **(A)** Schematic representation of the viral injections and experimental timeline. **(B)** Representative example of FACS cell sorting report showing the viable cells sorted according to Deep Red fluorescence and the NAc- and VTA-projecting mPFC neurons sorted according to mCherry and EGFP expression, respectively. **(C-D)** Relative mRNA expression of genes coding for receptors and proteins involved in cytoarchitecture and spine scaffolding in **(C)** NAc-projecting mPFC neurons and **(D)** VTA-projecting mPFC neurons. Significance was determined using two-tailed independent sample t-tests. Symbols and bars represent the group average ± SEM; * *p* < 0.05, ** *p* < 0.01, *** *p* < 0.005.

Overall, our findings are consistent with previous reports in human MDD post-mortem tissue and mouse models of chronic stress (42,45,47,50) and further suggest that CVS interferes with the activity of transcriptional programs controlling neuronal plasticity in a sex- and pathway-specific fashion.

### Opposite effects of activation and inhibition of the cortico-accumbal pathway on stress response

Our results suggest that CVS induces pathway-specific changes in the functional properties of NAc- and VTA-projecting mPFC neurons. We next tested whether reproducing these changes in stress-naïve mice was sufficient to trigger the expression of stress responses in a sex-specific fashion. To do so, we performed chemogenetic manipulations using our inter-sectional viral approach to express Designer Receptors Exclusively Activated by Designer Drug (DREADD; excitatory hM3Dq or inhibitory hM4Di; **Figures 6A-B**), and oriented our experiments toward the NAc-projecting mPFC neurons showing higher sEPSCs frequency and amplitude in females specifically (**Figures 3D-E**). First, we confirmed *in vitro* the efficacy of hM3Dq and hM4Di (**Supplemental Figure S3A-B**) to respectively stimulate and inhibit NAc-projecting mPFC neuron activity, which was similar for males and females. We then tested whether the acute activation of the cortico-accumbal pathway in stress naïve mice was sufficient to trigger depressive-like behaviors in both sexes (**Supplemental Figure S4A**). Our analysis shows that an acute stimulation of the cortico-accumbal pathway is not sufficient to induce the expression of anxiety, behavioral despair, and anhedonic-like behaviors in stress naïve males and females (**Supplemental Figures S4B-F**).

**Figure 6.**
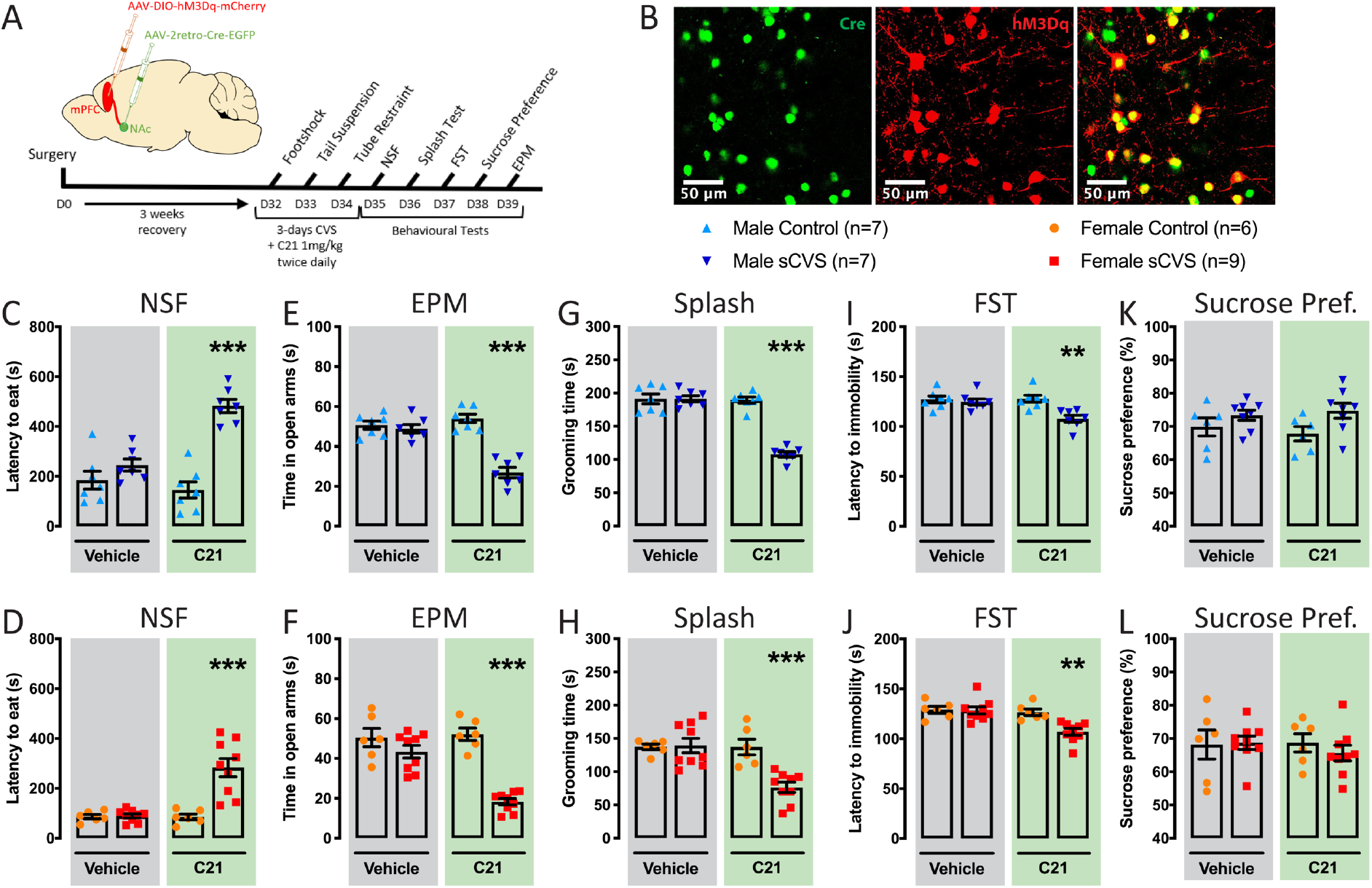
Sub-chronic variable stress (sCVS) coupled with overactivation of the NAc-projecting mPFC neurons induces anxiety and behavioral despair in males and females. **(A)** Schematic representation of the viral injections and experimental timeline. **(B)** Representative example of a viral infection in the mPFC with Cre in green and hM3Dq in red. **(C-J)** Chemogenetic activation of the NAc-projecting mPFC neurons during sCVS in males (blue) and females (red) induces anxiety and behavioral despair as illustrated in the **(C-D)** NSF, **(E-F)** EPM, **(G-H)** splash test and in the **(I-J)** FST. **(K-L)** Chemogenetic activation of the NAc-projecting mPFC neurons during sCVS fails to induce anhedonia in males (blue) and females (red). Significance was determined using two-way ANOVA with posthoc Tukey multiple comparisons tests. Symbols and bars represent the group average ± SEM; ** *p* < 0.01, *** *p* < 0.005.

We next tested whether cortico-accumbal hyperactivation during 3 days of variable stress (sCVS), a stress regimen insufficient to induce depressive-like behaviors (51), would trigger the expression of a stress response in a sex-specific fashion. Our results show that combining cortico-accumbal hyperactivity with sCVS for 3 days is sufficient to induce the expression of a stress response characterized by increased latency to eat in the NSF (males: F_(1,24)_=10.86, *p*=0.0030; females: F_(1,26)_=16.12, *p*=0.0004), decreased time spent in the open arms of the EPM (males: F_(1,24)_=17.57, *p*=0.0003; females: F_(1,26)_=14.18, *p*=0.0009), decreased time spent grooming in the splash test (males: F_(1,24)_=63.75, *p*<0.0001; females: F_(1,26)_=10.95, *p*=0.0027) and decreased the latency to immobility in the FST (males: F_(1,24)_=5.785, *p*=0.0242; females: F_(1,26)_=10.93, *p*=0.0028, **Figures 6C-J**) in both sexes compared to control and vehicle-treated groups. However, our analyses reported no group differences in the sucrose preference test (**Figures 6K-L**) in males and females compared to control groups, suggesting that hyperactivation of the NAc-projecting mPFC neurons during stress is sufficient for the expression of anxiety and behavioral despair but not anhedonia in males and females.

We then tested whether inhibition of the cortico-accumbal pathway would rescue the behavioral impact of CVS in males and females. To do so, we performed chemogenetic inhibition using our inter-sectional viral approach to express inhibitory DREADDs (hM4Di) in NAc-projecting mPFC neurons, in mice following 21 days of CVS. Behavioral assessment was performed in combination with acute injections of either C21 or vehicle to inhibit the activity of the cortico-accumbal pathway in males and females (**Figures 7A-B**). Interestingly, our results show clear sexually dimorphic behavioral responses to these stimulation patterns. As expected, 21 days of stress induced the expression of anxiety, behavioral despair, and anhedonia in both males and females (**Figures 7C-L**). However, the chemogenetic inhibition of NAc-projecting mPFC neurons through C21 administration failed to reverse the behavioral phenotype induced by 21 days of CVS in stressed males compared to controls (**Figure 7C-G**). In contrast, inhibition of the cortico-accumbal pathway following 21 days of CVS in females reversed the effects of stress in the NSF (F_(1,28)_=36.79, *p*<0.0001), EPM (F_(1,28)_=2752, *p*<0.0001), splash tests (F_(1,28)_=8.803, *p*=0.0061) and FST (F_(1,28)_=5.922, *p*=0.0216, **Figure 7H-K**) compared to female control groups. Interestingly, this stimulation paradigm failed to normalize the impact of CVS on the sucrose preference (**Figure 7L**) tests, consistent with the lack of effect observed on this behavioral parameter following the combined sub-chronic CVS and chemogenetic stimulation of this pathway in females (**Figures 6L**).

**Figure 7.**
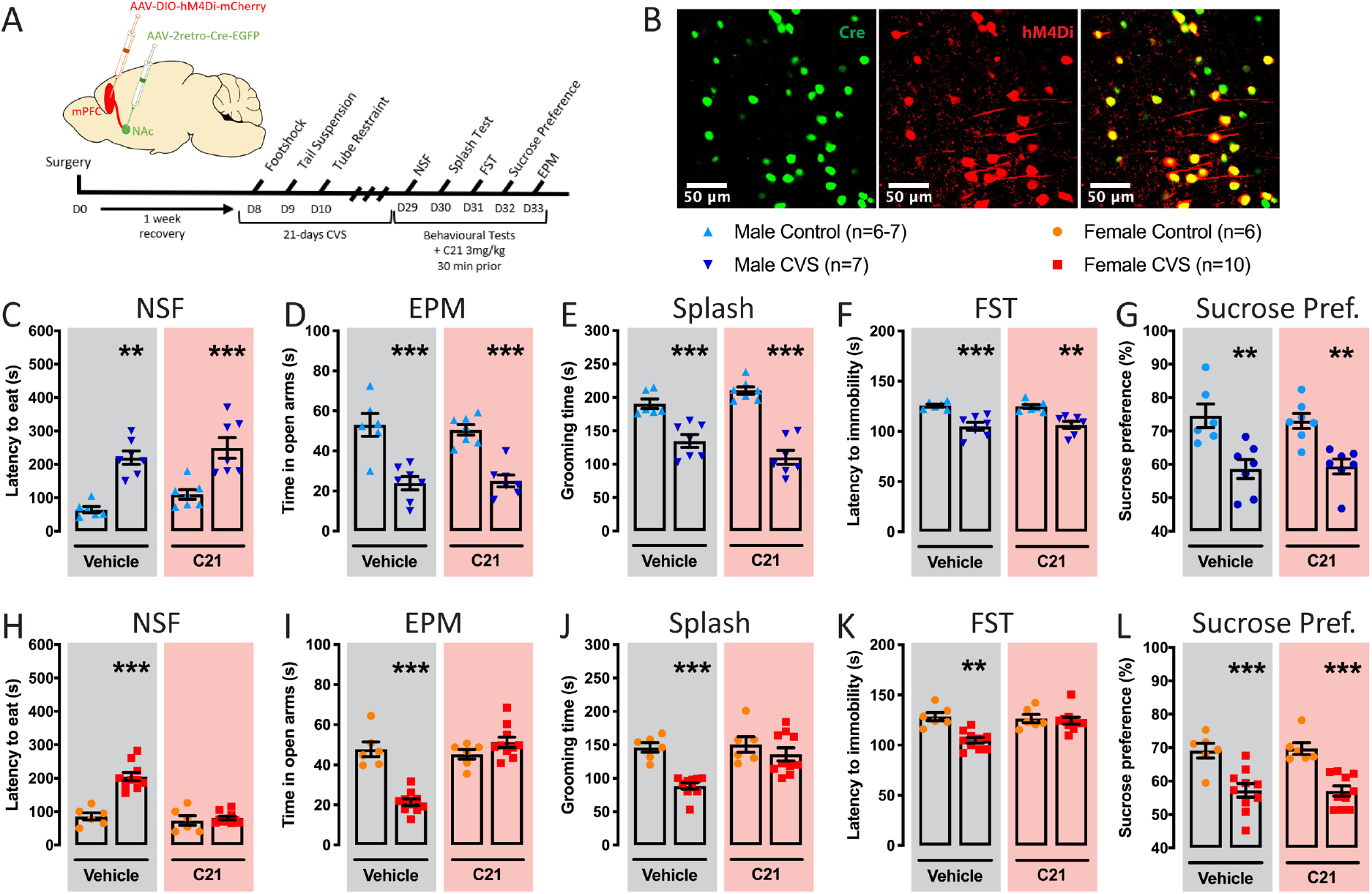
Inhibition of the NAc-projecting mPFC neurons rescues anxiety and behavioral despair induced by CVS in females only. **(A)** Schematic representation of the viral injections and experimental timeline. **(B)** Representative example of a viral infection in the mPFC with Cre in green and hM4Di in red. **(C-G)** Inhibition of the NAc-projecting mPFC neurons after CVS in males fails to rescue anxiety-like behavior in males as illustrated in the **(C)** NSF and **(D)** EPM, behavioral despair in the **(E)** splash test and **(F)** FST, and (**G**) anhedonia in the sucrose preference test. **(H-K)** Inhibition of the NAc-projecting mPFC neurons after CVS in females rescues anxiety and behavioral despair as illustrated by the **(H)** NSF, **(I)** EPM, **(J)** splash test and **(K)** FST. **(L)** Inhibition of the NAc-projecting mPFC neurons after CVS fails to rescue anhedonia in females. Significance was determined using two-way ANOVA with posthoc Tukey multiple comparisons tests. Symbols and bars represent the group average ± SEM; ** *p* < 0.01, *** *p* < 0.005.

Together, these results suggest that overactivation of the cortico-accumbal pathway in males and females is sufficient to induce the expression of anxiety and behavioral despair in males and females, but its inhibition reverses these behavioral phenotypes in females only. Interestingly, our results also suggest that NAc-projecting mPFC neurons exert a poor control, if any, over the expression of anhedonic behaviors in males and females.

## Discussion

Through its cortico-limbic projections, the mPFC controls the processing of emotional cues and modulate the expression and consolidation of stress responses (52,53). Here, we provide a comprehensive assessment of how chronic stress impacts the functional, morphological, and molecular properties of NAc- and VTA-projecting mPFC neurons in males and females highlighting the contribution of the cortico-accumbal pathway to the expression of stress responses in a sex-specific fashion.

Our results suggest that CVS induces a sex-specific reorganization of the cortico-accumbal and cortico-tegmental pathways’ dendritic and spine architecture. Importantly, global dendritic retraction has been consistently reported in the mPFC of human MDD (6,7,54) and different rodent models of chronic stress (8,9). Such morphological changes have been usually associated with either a reduction (10,55) or no change (56,57) in spine density. Interestingly, by investigating NAc- and VTA-projecting mPFC neurons independently, not only did we confirm the stress-induced attrition of apical and basal dendrites as previously observed (8,9), we also associated these changes to specific neuronal populations projecting to different limbic structures, demonstrating that NAc-projecting mPFC neurons are more severely affected in females and VTA-projecting mPFC neurons in males. Furthermore, by sorting out both neuronal populations, we provided insights into the molecular mechanisms underlying these effects in a sex-specific fashion. Indeed, *Dlg4* and *Shank1* encode for scaffolding proteins involved in spine maturation (41,58) and *Homer1* is required for the role of *Shank1* in excitatory spine maturation (59). Importantly, changes in the expression of these genes have been previously reported in the mPFC of human MDD and mouse models of chronic stress (42,46,48,49) although sex and cellular specificity was unknown. Together, our results suggest that by changing the expression of these genes in a sex- and pathway-specific fashion, CVS induces a global reorganization of NAc- and VTA-projecting mPFC neurons interfering with the functional activity of these neuronal circuits controlling stress responses in males and females.

Previous studies assessing the activity of the mPFC following chronic stress have yielded contrasting results with evidence in humans and mouse models supporting either elevated (11,60) or decreased (61,62) activity. For instance, deep brain stimulation in the mPFC was shown to induce anti-depressant-like responses in rats predisposed to helplessness (63). Similar effects were observed following the optogenetic stimulation of either Drd1-expressing mPFC neurons (64) or the whole mPFC (65) promoting resilience to chronic social defeat stress. In contrast, the optogenetic activation of NAc-projecting mPFC neurons was shown to modulate reward processing and seeking (29,40), decrease preference for social interaction in mice (14) and suppress natural reward-motivated behaviors in rats (15). Our findings suggest that the activity of NAc- and VTA-projecting mPFC neurons is potentiated by CVS by shifting the ratio of excitatory/inhibitory inputs toward excitation in a sex-specific fashion. The mPFC is a region integrating inputs from several brain regions (66,67), the activity of which has often been reported to be potentiated by chronic stress (30,61,68,69). For instance, activation of the amygdala (68) and hippocampal (69,70) afferences to the mPFC with optogenetic or chemogenetic was shown to promote anxiety in males while their inhibition reverses these effects (69). Importantly, inputs to the mPFC are modulated by GABAergic neurons, including somatostatin (SST) neurons controlling the spiking inputs to pyramidal neurons and parvalbumin (PV) cells regulating the spiking outputs from pyramidal neurons to projecting brain areas (71,72). Accordingly, molecular and functional modifications in mPFC somatostatin and parvalbumin neurons have been associated with the expression of stress responses (73,74), and their inhibition was shown to promote depressive-like behaviors in mice (35). Here, using our trans-sectional viral approach, we provide evidence suggesting that CVS induces a pathway-specific dysregulation of inhibitory and excitatory inputs affecting NAc-projecting mPFC neurons more severely in females and VTA-projecting mPFC neurons in males and females more evenly. More work will be required to understand the impact of chronic stress on the function of GABAergic interneurons in gating the inputs to NAc and VTA-projecting mPFC neurons and to determine the contribution of the different neuronal afferences to the mPFC from projecting brain structures in stressed males and females.

Stress response is defined by a vast spectrum of complex behavioral alterations affecting different behavioral domains relevant to the clinical manifestations of MDD in humans. Here, we present evidence supporting the contribution of the cortico-accumbal pathway in controlling the expression of behavioral despair and anxiety-like behaviors in males and females. Considering our findings, one could hypothesize that the reduction of inhibitory inputs on NAc-projecting mPFC neurons, as observed in males and females, primes the circuit toward hyperactivity, but not enough to trigger abnormal stress responses. Now, when combined with increased excitatory tones on these neurons, the shift toward excitation induced by CVS may become sufficient to trigger the expression of behavioral despair and anxiety-like behaviors as observed in stressed females. Consistently, once primed by sub-threshold stress, chemogenetic hyperactivation of NAc-projecting mPFC neurons triggers anxiety and depressive-like behaviors in both sexes while inhibition of this pathway rescues these behavioral effects but only when the pathway is pre-activated as observed in females. This being said, CVS induces despair and anxiety in males as well suggesting that redundant circuits may trigger the expression of these behavioral alterations. Consistently, previous studies using optogenetic approaches showed that anxiety-like behaviors in males are controlled by other circuits involving inputs to the mPFC from the basolateral amygdala (68) and ventral hippocampus (69,70). Besides, the stimulation of basomedial amygdala-projecting mPFC neurons was also shown to repress anxiety-like behaviors (75). Together, these findings provide important clinical insights into the neuronal substrate underlying the expression of behavioral alterations by chronic stress in males and females. Future work should determine whether different timing, duration and intensity of stimulation could be used before considering these neuronal targets for the treatment of depressive-like behaviors in a sex-specific fashion.

While consistent with some recent findings (11,68,69,76,77), our results are also in opposition with the conclusions from previous studies (20,53,65). These discrepancies can be explained in part by the different methodological approaches used in these studies including neuronal specificity and stimulation patterns. For instance, stimulation patterns (i.e. frequency and duration) used in optogenetic studies, with exceptions (15), are believed to go beyond natural pattern of neuronal activity (78). In contrast, the chemogenetic approach used in our study allowed to reproduce more physiologically relevant neuronal patterns and to dissect the contribution of the cortico-accumbal pathway in stress responses in a sex-specific fashion.

To conclude, the results presented here provide solid evidence uncovering the impact of chronic stress on the functional, morphological and molecular properties of two of the most important outputs from the mPFC. It will be important for future work to test whether different chronic stress paradigms impact the activity of NAc- and VTA-projecting mPFC neurons similarly. Indeed, while distinct mouse models have been shown to reproduce common and distinct transcriptional alterations relevant to MDD (13), it will be important to determine whether these clinically relevant molecular pathways result in similar functional impacts in males and females. Nevertheless, by revealing how stress impacts NAc- and VTA-projecting mPFC neurons in males and females, our findings identify these two cellular populations as neuronal substrates for the treatment of specific clinical manifestations of MDD in men and women.

## Acknowledgments

We would like to thank Drs. Annie Barbeau and Khaled Abdallah for their help with the experiments. We also thank the Plateforme d’Outils Moléculaires and the Plateforme de Tri Cellulaire (https://neurophotonics.ca/fr/pom) at CERVO Brain Research Centre for viral vector production and FACS sorting. BL holds a Sentinelle Nord Research Chair, is supported by a NARSAD Young Investigator Award, the Canadian Institutes of Health Research (Grant No. SVB397205), and the Natural Science and Engineering Research Council of Canada (Grant No. RGPIN-2019-06496), and receives Fonds de Recherche en Santé du Québec (FRQS) Junior-1 salary support. CFS lab was supported by NSERC grants RGPIN-2020-06376 and DGECR-2020-00060. CDP lab was supported by CIHR grant PJT169117 and NSERC grant RGPIN-2017-06131. IT lab was supported by CIHR grant MOP136967.

## Author contributions

T.P.B. and B.L. conceived the project, designed the experiments, and wrote the manuscript. T.P.B. also generated and analyzed all the data. M.C.P. and C.F.S. contributed to the morphological analyses. J.C.H.S., J.S., I.T., and C.D.P. generated the electrophysiological data. F.Q. and A.A.L. helped with *in vivo* surgeries. M.C.P., D.M., and E.A. contributed to behavioral experiments. L.J.B contributed to the molecular experiments. Y.C. generated the DREADD AAVs. All authors contributed to the preparation of the manuscript.

## Supplemental Information

### Detailed Methods

#### Animals

All animal experiments were carried out in male and female C57BL/6NCrl mice (Charles River Laboratories, Kingston, NY) aged from 8 to 14 weeks old. Experiments were performed following the Canadian Guide for the Care and Use of Laboratory Animals and were approved by the Animal Protection Committee of Université Laval. All animals were housed in standard conditions (4 per cage, at 22±1°C, on a 12h light cycle, lights on at 8 am) without any enrichment. Food and water were provided ad libitum unless specified otherwise.

#### Chronic and Subchronic Variable Stress Paradigm

Depressive-like behaviors in males and females were induced through chronic variable stress (CVS) as described before (1,2). Briefly, CVS consists of three different stressors repeated for 21 days after which both males and females exhibit a range of behavioral alterations including anhedonia, behavioral despair, and anxiety. On the first day, mice are placed in a shocker in which 100 footshocks at 0.45mA are applied randomly in one hour. On day two, mice are suspended by the tail for one hour. Finally, on the third day, mice are restrained in a 50mL falcon tube for one hour. These stressors are repeated once per day over 21 days. For both males and females, a group of unstressed control mice was gently handled 2 minutes per day.

A subthreshold version of CVS (sCVS) was used to study stress susceptibility (1). This duration of stress, by itself, is insufficient to induce stress-related behavioral abnormalities in males and females, but when combined with molecular or physiological challenges, can yield enhanced stress susceptibility _(2,3)_. The same stressors described above were repeated for 3 days 3 weeks after surgeries in males and females. The following day, the behavior was assessed using the behavioral assays described below.

#### Behavioral Assessment

A panel of behavioral tests was used to assess anxiety-, behavioral despair- and anhedonia-like states in male and female mice as previously described (1,2). At the end of the CVS/sCVS paradigms, mice were single-housed and behavioral tests were performed once per day in the following order: novelty-suppressed feeding test, splash test, forced swim test, sucrose preference, and elevated-plus maze test. All behavioral assessments were performed in the morning, in a dedicated room, different from the housing and stress rooms. On each testing day, mice were habituated to the test room one hour before testing and transferred back to the housing room immediately after testing.

##### Novelty-Suppressed Feeding Test

The behavioral test was adapted from a published protocol (1,3,4). Mice were food-deprived for 24h prior to testing. Mice were placed in an open field box of 50 x 50 x 40 cm with corn cob bedding and a single food pellet in the middle under red-light conditions. Mice were allowed to explore for a maximum of 10 minutes or until they started to eat. The latency to eat was scored as a measure of anxiety. Mice were then immediately transferred to their home cage containing a single food pellet and the latency to eat was scored for up to 3 minutes under standard light conditions.

##### Splash Test

The behavioral test was adapted from a published protocol (1,3,5). Mice were sprayed on the back with a 10% sucrose solution three times and then placed into an empty housing cage under red light conditions. The behavior was recorded for 5 min via videotape. The total amount of time grooming over the 5 min period was hand-scored by an observer blind to experimental conditions. Grooming activity included nose/face grooming, head, and body washing.

##### Forced Swim Test

The behavioral test was adapted from a published protocol (1,3). The forced swim test (FST) was performed under white light conditions. Mice were placed in a 3L beaker containing 2L of water (25±1°C). Mice activity was recorded for a total of 6 minutes using an automated system (ANYmaze, version 5, Stoelting Co., Wood Dale, IL). Immobility was defined as the absence of struggling with minimal activity to maintain head above water. The latency to immobility was assessed.

##### Sucrose Preference Test

The behavioral test was adapted from a published protocol (1,3). Mice were given two bottles filled with water for a 24 hours habituation period. The following day, one of the two bottles was replaced with a 1% sucrose bottle for 24 h. The two bottles were then weighed and the position was switched for an additional 24 h. The total duration of the test was 48 h. Sucrose preference was calculated by determining the percentage of total sucrose consumption divided by total liquid consumption (sucrose + water).

##### Elevated-Plus Maze Test

The behavioral test was adapted from a published protocol (2). Mice were placed in the middle of an elevated-plus maze apparatus under red light conditions. Each arm of the maze measured 50 x 10 cm. The black Plexiglas cross-shaped maze consisted of two open arms with no walls and two closed arms (40 cm high walls) and was on a pedestal 50 cm above floor level. Behavior was tracked using an automated system (ANYmaze, version 5). Behavior was measured as total time spent in combined open arms and total time in combined closed arms.

##### Emotionality score

The emotionality score for each mouse was calculated as previously described (6). An individual z-score was calculated for each test using the following formula:

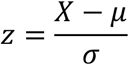

in which X represents the individual value, µ the control group mean, and σ the control group standard deviation. Decreasing parameters such as time spent in the open arms (EPM), grooming time (splash test), and sucrose consumption (sucrose preference test) were converted into positive changes compared to group means to indicate increased emotionality. Same sex controls were used to compare the data. An emotionality score was then calculated by averaging the z-scores obtained for each parameter.

#### Estrous cycle

The female estrous cycle was monitored in the female mice by vaginal smears during two consecutive days for more accuracy, either at the beginning of the behavioral tests or before being euthanized if no behavioral test was performed. Briefly, 20µL of sterile saline 0.9% was flushed in the vagina 2-3 times and then put on a glass slide. The slides were observed without any staining under a light microscope with a 10x objective. The smears were classified into four categories as previously described (7): proestrus, estrus, metestrus, and diestrus.

#### Viral vectors and stereotaxic injections

Adeno-associated viruses (AAVs) used in this study along with providers and titers are listed in **Supplemental Table 1**.

pAAV-hSyn-Cre and pAAV-CAG-Cre-EGFP-WPRE were developed by the POM. pAAV-EF1a-Flpo and pAAV-Ef1a-fDIO mCherry were a gift from Karl Deisseroth. pAAV-hSyn-DIO-EGFP, pAAV-hSyn-DIO-mCherry, pAAV-hSyn-DIO-hM3D(Gq)-mCherry and pAAV-hSyn-DIO-hM4Di-mCherry were a gift from Bryan Roth.

AAV-hSyn-DIO-hM3Dq-mCherry and AAV-hSyn-DIO-hM4Di-mCherry vectors from serotypes 2/9 were obtained as described earlier (8). Briefly, the serotype 2/9 AAVs were generated by tripartite transfection (pAAV-rep2/cap9 capsid plasmid, adenovirus helper plasmid, and AAV-vector plasmid) into HEK293-derived AAV-293 cells (Stratagene, #240073). Three days after transfection, the virus was extracted and then quantified by quantitative polymerase chain reaction.

AAVs were injected in the mPFC, the NAc, and/or the VTA of 8 weeks old C57BL/6 mice. Mice were anesthetized with isoflurane and placed on a stereotaxic frame (Stoelting Co., Wood Dale, IL). Then, viral vectors were injected bilaterally at a rate of 1µL/min followed by 5 min of rest. The stereotaxic coordinates used and injected volumes are reported in **Supplemental Table 2**.

#### Electrophysiological recordings

Mice were habituated one week to the animal facility prior to any manipulations. In total, 16 male and 15 female mice were injected with AAV2-retro-EF1a-FlpO in the NAc, AAV2-retro-hSyn-Cre in the VTA, and a 1/1 ratio of AAV-EF1a-fDIO-mCherry/AAV-hSyn-DIO-EGFP in the mPFC. One week after surgery, 8 male and 7 female mice underwent CVS for 21 days and 8 males and females were handled once a day for 2 minutes as control. After 21 days, we obtained acute brain slices using a vibratome (VT2000; Leica). Both stressed and control mice were anesthetized with isoflurane and transcardially perfused with 10 ml of an ice-cold NMDG-artificial cerebrospinal fluid (aCSF) solution containing (in mM): 1.25 NaH_2_PO_4_, 2.5 KCl, 10 MgCl_2_, 20 HEPES, 0.5 CaCl_2_, 24 NaHCO_3_, 8 D-glucose, 5 L-ascorbate, 3 Na-Pyruvate, 2 Thiourea, 93 NMDG (osmolarity adjusted with sucrose to 300–310 mOsmol/l) and pH is ajusted to 7.4 with HCl 10N. Kynurenic acid (2 mM) was added to the perfusion solution on the day of the experiment. Brains were then quickly removed and placed in the cutting chamber filled with ice-cold perfusion solution (0-4°C) and sliced at 250 µm. Slices were first placed in a 32°C oxygenated perfusion solution for 10 minutes, before being incubated for one hour at room temperature in HEPES-aCSF solution (in mM: 1.25 NaH_2_PO_4_, 2.5 KCl, 10 MgCl_2_, 20 HEPES, 0.5 CaCl_2_, 24 NaHCO_3_, 2.5 D-glucose, 5 L-ascorbate, 1 Na-Pyruvate, 2 Thiourea, 92 NaCl, 20 Sucrose (osmolarity adjusted to 300–310 mOsmol/l at pH 7.4) and finally transferred into a recording chamber on the stage of an upright microscope (Zeiss) where it was perfused at a rate of 3–4 ml/min with artificial cerebrospinal solution (aCSF in mM: 120 NaCl, 5 HEPES, 2.5 KCl, 1.2 NaH_2_P0_4_, 2 MgCl_2_, 2 CaCl_2_, 2.5 glucose, 24 NaHCO_3_, 7.5 sucrose). The perfusion chamber and the aCSF were kept at 32°C. All solutions were oxygenated at 95% O_2_/5% CO_2_.

A water immersion x60 objective and a video camera (Zeiss, Germany) were used to visualize the glutamatergic pyramidal neurons in the mPFC expressing either mCherry in layer II/III of the prelimbic cortex (NAc-projecting mPFC neurons) or EGFP in layer V of the prelimbic cortex (VTA-projecting mPFC neurons), and blue light was delivered through the objective with a Colibri 7 LED light source (Zeiss). Whole-cell voltage-clamp recordings were performed with an Axopatch 200B amplifier (Molecular Devices, San Jose, CA) using borosilicate patch electrodes (3-7 MΩ resistance). Pipettes were filled with an intracellular patch solution containing (in mmol/l): 115 Cesium methanesulfonate, 20 Cesium chloride, 10 HEPES, 2.5 MgCl2, 4 Na2-ATP, 0.4 Na-GTP, 10 Na-phosphocreatine, 0.6 EGTA, 5 QX314 and 0.2% Biocytin (pH 7.35).

Signals were filtered at 5 kHz using a Digidata 1500A data acquisition interface (Molecular Devices, San Jose, CA) and acquired using pClamp 10.6 software (Molecular Devices, San Jose, CA). Pipette and cell capacitances were fully compensated. Passive membrane properties were measured. Spontaneous excitatory postsynaptic currents (sEPSCs) were recorded at −60 mV, and spontaneous inhibitory postsynaptic currents (sIPSCs) at 0 mV.

A total of 182 neurons in 31 mice was recorded. For the neurons where both sEPCSs and sIPSCs were recorded, a frequency ratio of sEPSCs/sIPSCs was calculated. A total of 56 NAc-projecting and 62 VTA-projecting mPFC neurons were used in the sE/IPSC frequency ratio. Variations in passive membrane properties, sE/IPSC frequency ratio, and amplitude were assessed using two-way ANOVA with stress and sex as main factors and posthoc Tukey tests in NAc- and VTA-projecting mPFC neurons separately. Significance was fixed at p<0.05.

#### Morphological analyses

Mice were habituated one week to the animal facility prior to any manipulations. We used mice aged 8 weeks and injected them stereotaxically with AAV2-retro-hSyn-Cre in either NAc or VTA diluted at 1:10^4^ and the AAV-hSyn-DIO-EGFP in the mPFC. In total, 12 males and 12 females were injected. One week after surgery, 6 males and 6 females mice underwent CVS for 21 days and 6 males and 6 females were handled once a day for 2 minutes as control. After 21 days, both stressed and control mice were deeply anesthetized using a 20% urethane solution injected intraperitoneally. Residual blood was washed out with intracardiac perfusion with ice-cold saline (0.9%) followed by filtered ice-cold paraformaldehyde (PFA 4%). Brains were collected and post-fixed in PFA overnight and then cryoprotected in 30% sucrose solution for at least 48 hours. The brains were cut in 100µm-thick slices using a microtome (Microm HM430, ThermoFisher) and placed in PBS. The slices were washed 3 times for 5 minutes in PBS before being mounted on glass slides. After 2 hours of drying in the dark, the slides were mounted with a protecting medium (ProLong Gold Antifade Reagent, Invitrogen, #P36930) and coverslipped.

Picture acquisition was performed with a confocal microscope (Nikon A1R HD) with a 60x oil-immersion objective. Stacked images of the labeled glutamatergic pyramidal neurons in the mPFC (encompassing prelimbic cortex) were captured with 0.5µm between each picture. Morphological 3D reconstruction of the neurons and spine detection was performed using Neurolucida 360 (MBF Bioscience, Williston, VT). Scholl analysis was performed starting at 10µm from the soma with 5µm intervals as previously described (9). Automatic spine detection was used to identify spines on the full length of the apical dendrite. Spine automatic detection was supervised and manually corrected by a blind investigator. Three mice per group (n=3) with 9-10 neurons per mouse were used for the analyses. Variations in the number of intersections were assessed using two-tailed independent sample t-tests. Variations in spine density were assessed using two-way ANOVA with stress and sex as main factors and Fisher’s LSD posthoc tests in NAc- and VTA-projecting mPFC neurons separately. Significance was fixed at p<0.05.

#### Molecular experiments

Mice were habituated one week to the animal facility prior to any manipulations. In total, 12 male and 12 female mice were injected with AAV2-retro-EF1a-FlpO in the NAc, AAV2-retro-hSyn-Cre in the VTA, and a 1:1 ratio of AAV-EF1a-fDIO-mCherry/AAV-hSyn-DIO-EGFP in the mPFC. One week after surgery, 6 males and 6 females underwent CVS for 21 days and 6 males and 6 females were handled once a day for 2 minutes as control. After 21 days, both stressed and control mice were sacrificed by cervical dislocation to collect mPFC tissue for fluorescence-assisted cell sorting (FACS) and qPCR using a 12-gauge punch on a 1mm-thick slice.

##### Neuron isolation

Brain tissue was processed for FACS as previously described (10). Briefly, the sample was placed on a rotator at 37°C in a 1mg/mL papain digestion buffer (5% w/v D-trehalose, 0.05 mM APV, 0.0125 mg/mL DNAse, in HibernateTM-A (Thermofisher, #A1247501)). After 45 minutes of incubation, the tissue was transferred into FACS buffer (5% w/v D-trehalose, 0.05 mM APV, 0.0125 mg/mL DNAse, in HibernateTM-A) and triturated with descending diameter of pipette tips. The dissociated cells were then passed through a 70µm filter, placed on an ovomucoid-albumin gradient solution (0.58mg/mL ovomucoid inhibitor-albumin (Worthington Biochemical, LK003182) in FACS buffer) and spun at 900RPM for 6min at 4°C. Pelleted cells were then resuspended in 400µL of FACS buffer. NAc- and VTA-projecting mPFC neurons were isolated based on their fluorescence (NAc: mCherry; VTA: EGFP). Neuronal cells were sorted on a FACSAriaII (BD Biosciences). MitoTracker Deep Red FM (ThermoFisher, #M22426) was used as a marker of viability. Neuronal cells from both pathways were collected in PBS and TRIzol-LS (1:3 ratio; ThermoFisher Scientific, #10296028).

##### RNA extraction and cDNA synthesis

Samples were then processed for RNA isolation with the Direct-Zol RNA MicroPrep kit (Zymo Research, #R2063) including a DNAse I digestion step. RNA was suspended in 12µL of nuclease-free water (Qiagen) at the end of extraction. RNA quantification and RNA integrity number (RIN) were determined on an Agilent 2100 Bioanalyzer (Agilent Technologies, Santa Clara, CA) using the Agilent RNA 6000 Pico Kit (#5067-1513).

Following quantification, RNA was reverse transcribed using the iScript cDNA synthesis kit (Bio-Rad, #1708891). 1 ng of RNA was used for NAc-projecting mPFC neuron samples and 0.5 ng for VTA-projecting mPFC neuron samples in a reaction volume of 20µL. Samples were diluted in nuclease-free water at a ratio of 1:10.

##### Real-time quantitative polymerase chain reaction

Primers were designed using the PrimerBlast tool (NCBI, Bethesda, MD) and are shown in **Supplemental Table 3**. RT-qPCR was performed with a QuantStudio 5 Real-Time PCR System (Applied Biosystems, Foster City, CA). The reactions were performed in MicroAmp Optical 384-Well Plates (Applied Biosystems, #4309849) in a 10µL volume with Luna Universal qPCR Master Mix (New England Biolabs, #M3003E). 2µL of cDNA was used per well with 2.5µM primers and each reaction was performed in triplicate. QuantStudio 5 Real-Time PCR System Software (Applied Biosystems) was used to analyze the data. For the determination of relative gene expression levels, the Comparative Ct method (11) was applied with GAPDH and 18S as reference genes. Relative gene expression levels were assessed with two-way ANOVA (with sex and stress as main factors) followed by Tukey posthoc tests. Significance was fixed at p<0.05.

#### Chemogenetic experiments

Chemogenetic manipulations were performed as described before for acute and chronic administration of DREADD agonist (12,13). To avoid any off-target effects of the usual DREADD actuator clozapine-N-oxide (14), we used DREADD Agonist 21 (Compound 21 (C21); Tocris, #6422) dissolved in sterile saline 0.9% (15,16).

##### *In vitro* validation of DREADDs efficiency

Mice were habituated one week to the animal facility prior to any manipulations. In total, 4 male and 4 female mice were injected with AAV-hSyn-DIO-hM3Dq-mCherry in the mPFC and AAV2-retro-CAG-Cre-EGFP-WPRE in the NAc, and 4 males and 4 females were injected with AAV-hSyn-DIO-hM4Di-mCherry in the mPFC and AAV2-retro-CAG-Cre-EGFP-WPRE in the NAc. Three weeks after surgery, mice were first anesthetized with ketamine-xylazine (100 and 10 mg/kg). The brain was then quickly dissected and maintained in ice-cold artificial cerebrospinal fluid (aCSF) containing the following: 124mM NaCl, 2.8mM KCl, 1.2mM CaCl_2_, 2 mM MgSO_4_, 1.25 mM NaH_2_PO_4_, 26 mM NaHCO_3_, and 10 mM D-glucose (Sigma-Aldrich Canada) (pH 7.4), aerated with 95% O_2_ and 5% CO_2_. Osmolarity was 300 ± 5 mOsm. Coronal slices (300-350 µm) were cut with a vibratome to obtain complete sections containing the infected region. Slices were transferred to a holding chamber where they were kept at room temperature for at least 1 hr in the same aCSF and aerated with 95% O_2_ and 5% CO_2_. The brain slices were transferred into a submerged recording chamber maintained at 34°C, containing the perfusion aCSF at a rate of 3 ml/min. The perfusion solution was identical to the cutting solution.

Infected region was identify using a CCD camera (PentaMax, Princeton Instrument) on a fluorescent upright microscope with optic filters for emission wavelengths of 568 nm. Neurons were preselected in the infected region using an infrared differential interference contrast camera microscopy on the same microscope. We obtained somatic whole-cell current-clamp recordings (10–20 MΩ access resistances) with patch pipettes (resistance between 3–5 MOhm) containing the following: 130 mM potassium D-gluconate, 10 mM 4-(2-hydroxyethyl)-1-piperazineethanesulfonic acid (HEPES), 10 mM KCl, 2 mM MgCl2, 2 mM ATP, 0.2 mM GTP (Sigma-Aldrich Canada) at pH 7.2 and 280 mOsm. 5-6 neurons per mouse were recorded at resting potential or a steady depolarizing current was injected to maintain the membrane potential near firing threshold. Stabilized recordings were obtained during 2 minutes before the bath solution was exchanged with the agonist 21 0.001% (C21)-containing aCSF solution for 3-5 minutes and then washed with aCSF solution.

##### Acute chemogenetic activation

Mice were habituated one week to the animal facility prior to any manipulations. In total, 28 male and 30 female mice were injected with AAV-hSyn-DIO-hM3Dq-mCherry in the mPFC and AAV2-retro-CAG-Cre-EGFP-WPRE in the NAc. Three weeks after surgery, 13 male and 13 female mice were subjected to a batterie of behavioral tests described above to assess depressive-like behaviors (NSF, splash test, FST, sucrose preference test, and EPM). 30 min prior to each test, 8 males and 8 females were administered C21 intraperitoneally at 3mg/kg. 5 males and 5 females were injected with sterile saline 0.9% as vehicle as controls. Behavioral consequences were assessed using a two-way ANOVA with sex and treatment as main factors and posthoc Tukey tests. Significance was fixed at p<0.05.

##### Sub-chronic chemogenetic activation

Male and female mice were divided into 4 groups: sCVS with C21, sCVS with saline, control with C21, and control with saline. In total, 14 males and 15 females underwent sCVS for 3 days, while 14 males and 15 females were gently handled for 2 min per day. On stress days, mice received either C21 or 0.9% saline intraperitoneal injections (1mg/kg), twice a day (8 am and 4 pm). Following sCVS, the batterie of behavioral tests described above was performed. Behavioral consequences were assessed using two-way ANOVA with stress and treatment as fixed effects and posthoc Tukey tests, separately in males and females. Significance was fixed at p<0.05.

##### Chemogenetic inhibition

Mice were habituated one week to the animal facility prior to any manipulations. In total, 27 male and 32 female mice were injected with AAV-hSyn-DIO-hM4Di-mCherry in the mPFC and AAV2-retro-CAG-Cre-EGFP-WPRE in the NAc. One week after surgery, 14 males and 16 females underwent a 21-day CVS protocol, while 13 males and 16 females were gently handled for 2 min per day. On the next day, the batterie of behavioral tests described above was performed. 30 min prior to each behavioral assessment, the mice were administered intraperitoneally 3mg/kg of C21 or sterile saline 0.9% as vehicle. Behavioral consequences were assessed using Two-way ANOVA with stress and treatment as main factors and posthoc Tukey tests, separately in males and females. Significance was fixed at p<0.05.

##### Tissue processing

At the end of the behavioral tests, mice were perfused as described earlier. Their brains were collected and cut in 40µm-thick slices with a microtome (Microm HM430, ThermoFisher). The slices were washed 3 times for 5 minutes in PBS before being mounted on glass slides. After 2 hours of drying in the dark, slides were mounted with a protecting medium (ProLong Gold antifade reagent, Invitrogen, #P36930) and coverslipped. Image acquisition was performed with a confocal microscope (Nikon A1R HD) with a 25x water-immersion objective to check the injection sites and spreading of the viruses. Four mice were classified as outliers because of a lack of infection and/or expression of the viruses in the mPFC.

##### Statistical analysis

Sample size calculation was not performed. However, we justified every experiment sample size based on several previously published reports using similar or even smaller sample sizes and showing the power to detect significant statistical differences.

All values given in the text and figures indicate mean ± standard error on the mean (SEM). *p*-values were considered significant if p < 0.05. On the figures, the stars are distributed as follow: * *p*<0.05, ** *p*<0.01 and *** *p*<0.005. All statistical analyses were achieved using GraphPad Prism 8 (GraphPad, San Diego, CA).

For <u>behavioral test analyses</u>, 32 mice were used (16 males: 8 stressed, 8 controls; 16 females: 8 stressed, 8 controls). Two-way ANOVA (with sex and stress as fixed effects) were performed, followed by posthoc Tukey tests for multiple comparisons.

For <u>electrophysiological analyses</u>, 31 mice were used (16 males: 8 stressed, 8 controls; 15 females: 7 stressed, 8 controls). In total, 87 NAc- and 95 VTA-projecting neurons were recorded for sEPSCs, and 75 NAc- and 85 VTA-projecting neurons for sIPSCs. Two-way ANOVA (with sex and stress as fixed effects) were performed with posthoc Tukey tests to assess the differences in frequency, amplitude, sE/IPCSc frequency ratio, membrane input resistance, and capacitance.

For <u>morphological analyses</u>, 24 mice were used (12 males: 3 NAc stressed, 3 NAc controls, 3 VTA stressed, 3 VTA controls; 12 females: 3 NAc stressed, 3 NAc controls, 3 VTA stressed, 3 VTA controls). Mixed-effects model for repeated measures (with stress or sex as fixed effect and distance as repeated measures) with posthoc Sidak multiple comparisons test was used to compare the number of intersections. Two-way ANOVA (with sex and stress as fixed effects) was used for total and mature spine density with posthoc Tukey tests for multiple comparisons.

For <u>molecular analyses</u>, 24 mice were used (12 males: 6 stressed, 6 controls; 12 females: 6 stressed, 6 controls). Two-way ANOVA (with sex and stress as fixed effects) were used to compare the relative expression of each gene with Gapdh and 18S as reference genes with posthoc Tukey tests for multiple comparisons.

For <u>chemogenetic acute activation analyses</u>, 26 mice were used (13 males: 8 C21, 5 vehicle; 13 females: 8 C21, 5 vehicle). Behavioral consequences were assessed with Two-way ANOVA (with sex and treatment as fixed effects), followed by posthoc Tukey tests for multiple comparisons.

For <u>chemogenetic sub-chronic activation analyses</u>, 60 mice were used (28 males: 7 C21 sCVS, 7 vehicle sCVS, 7 C21 controls, 7 vehicle controls; 34 females: 10 C21 sCVS, 10 vehicle sCVS, 7 C21 controls, 7 vehicle controls). 1 female C21 sCVS, 1 female C21 control, and 1 female vehicle control did not exhibit DREADD expression in the mPFC and were removed from the analyses. 1 female vehicle sCVS was an outlier in all the tests, as determined by the Grubbs test, and was removed from the analyses. Behavioral consequences were assessed with Two-way ANOVA (with stress and treatment as fixed effects), followed by posthoc Tukey tests for multiple comparisons, males, and females separately.

For <u>chemogenetic acute inhibition analyses</u>, 62 mice were used (28 males: 7 C21 CVS, 7 vehicle CVS, 7 C21 controls, 7 vehicle controls; 34 females: 10 C21 CVS, 10 vehicle CVS, 7 C21 controls, 7 vehicle controls). 1 female vehicle control and 1 female C21 control did not exhibit DREADD expression in the mPFC and were removed from the analyses, and 1 male vehicle control died from battle wounds before the beginning of the behavioral tests. Behavioral consequences were assessed using Two-way ANOVA (with stress and treatment as fixed effects), followed by posthoc Tukey tests for multiple comparisons, males, and females separately.

## Supplemental Figures

**Supplemental Figure S1.**
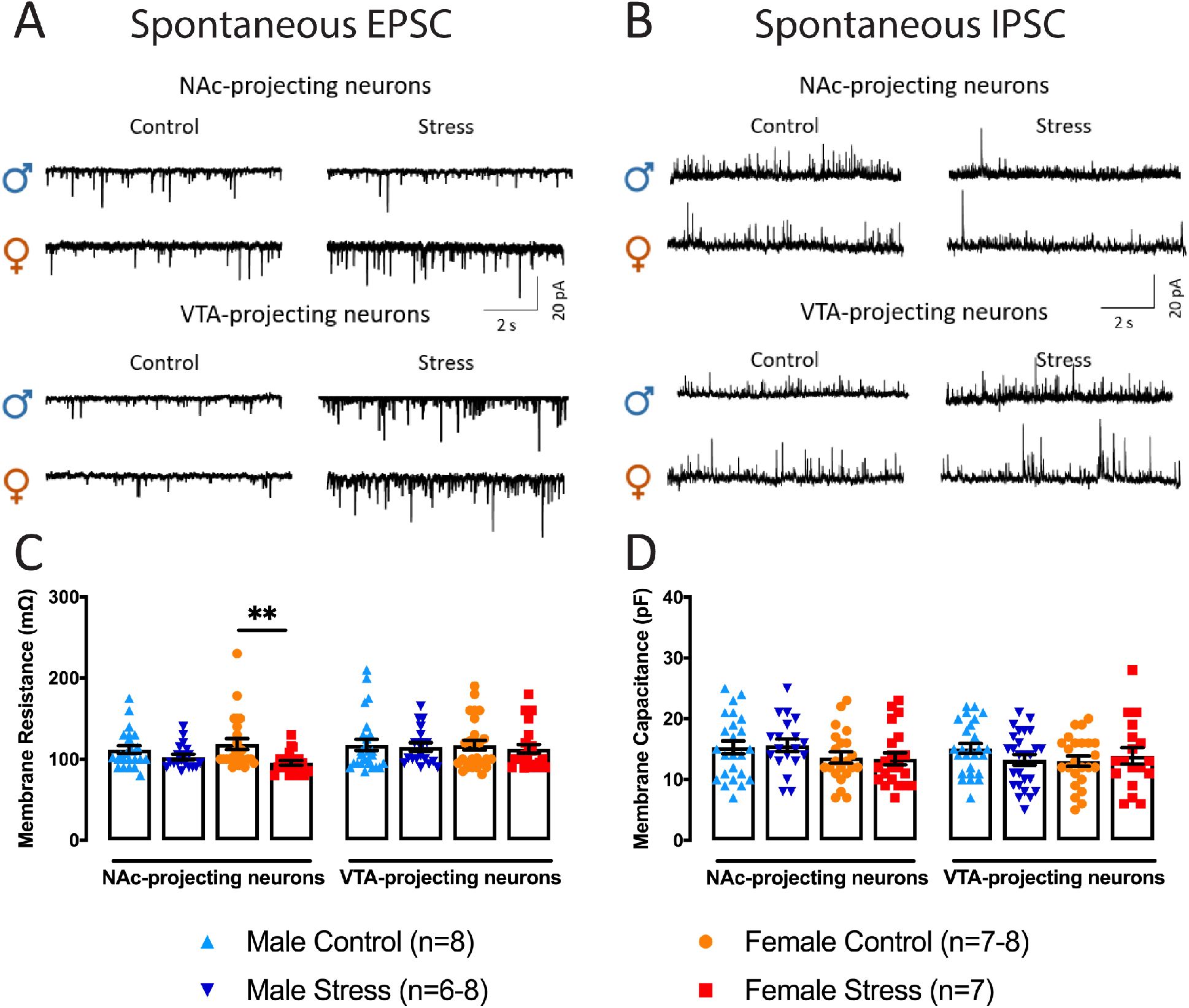
Passive membrane resistance is reduced in NAc-projecting mPFC neurons in stressed females only. **(A-B)** Representatives traces of spontaneous EPSCs **(A)** and spontaneous IPSCs **(B)** in both neuronal populations from males and females. **(C)** Passive membrane resistance is reduced only in NAc-projecting mPFC neurons in stressed females and not in stressed males. **(D)** No stress-induced changes in membrane capacitance were reported in NAc- and VTA-projecting mPFC neurons in either sex. Significance was determined using two-way ANOVA with *post-hoc* Tukey multiple comparisons test. Symbols and bars represent the group average ± SEM; ** *p* < 0.01.

**Supplemental Figure S2.**
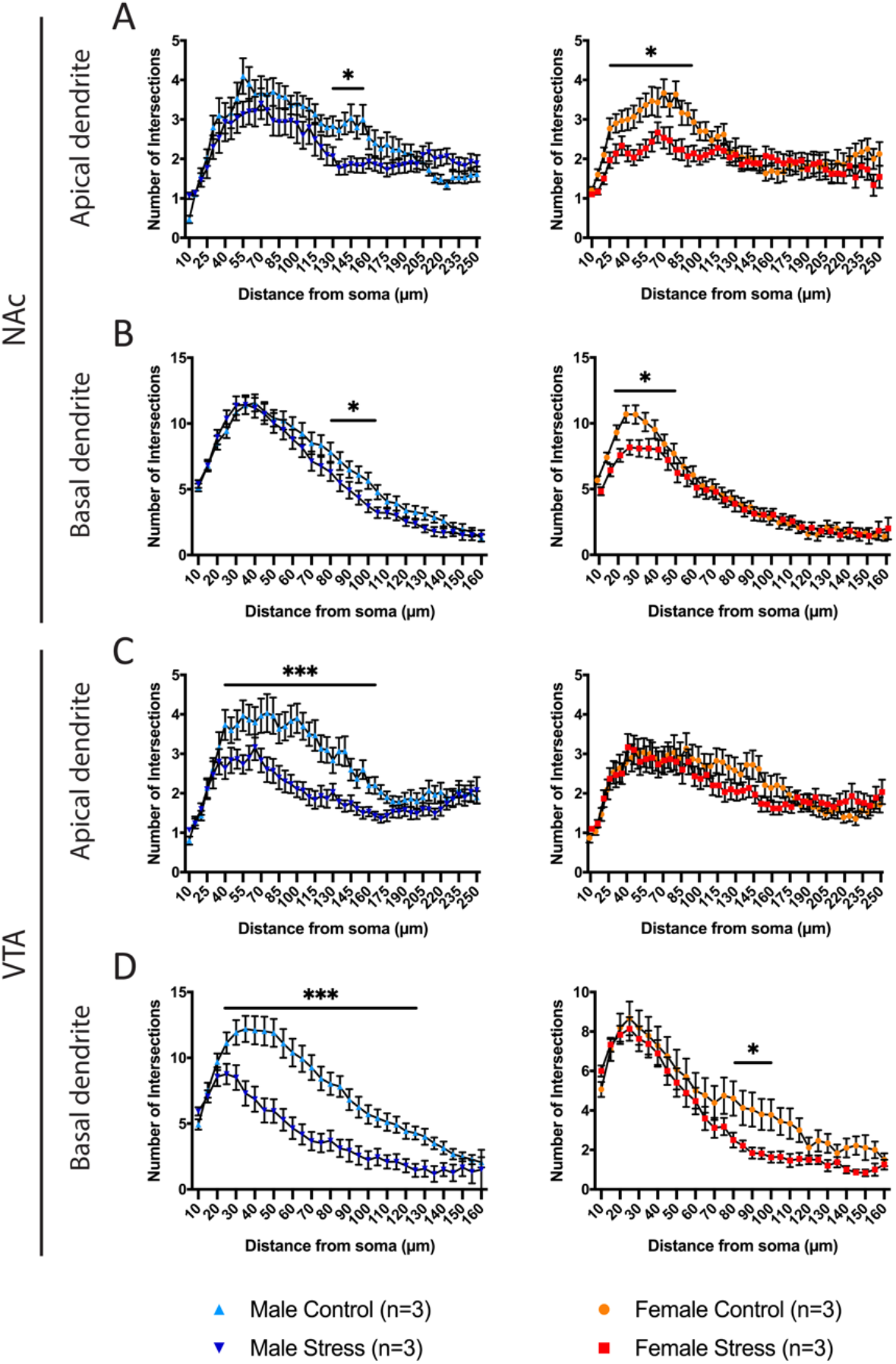
Detailed Sholl analysis of the apical and basal dendrites of NAc-projecting and VTA-projecting mPFC neurons after CVS in males and females. Sholl analyses of the number of intersections relative to the distance from the soma for apical **(A-C)** and basal **(B-D)** dendritic branches NAc-projecting and VTA-projecting mPFC neurons in males and females. Significance was determined using two-way ANOVA with *post-hoc* Tukey multiple comparisons test. Symbols and bars represent the group average ± SEM; * *p* < 0.05, ** *p* < 0.01, *** *p* < 0.005.

**Supplemental Figure S3.**
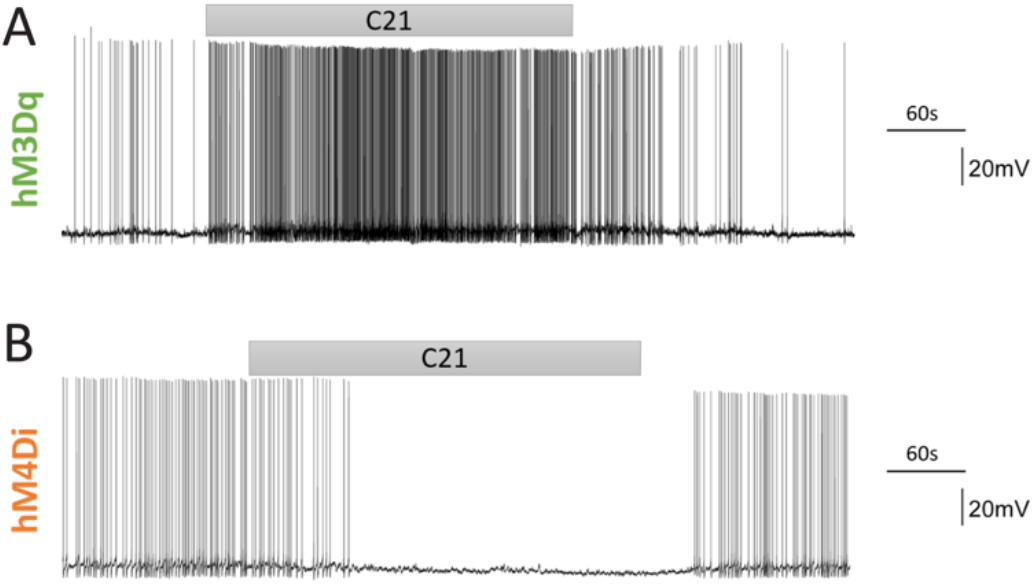
C21 is efficiently acting on DREADD-infected NAc-projecting mPFC neurons *in vitro*. **(A)** Application of C21 is efficiently increasing NAc-projecting mPFC neuron activity after infection with the excitatory DREADD hM3Dq. **(B)** Application of C21 is efficiently inhibiting NAc-projecting mPFC neuron activity after infection with the inhibitory DREADD hM4Di.

**Supplemental Figure S4.**
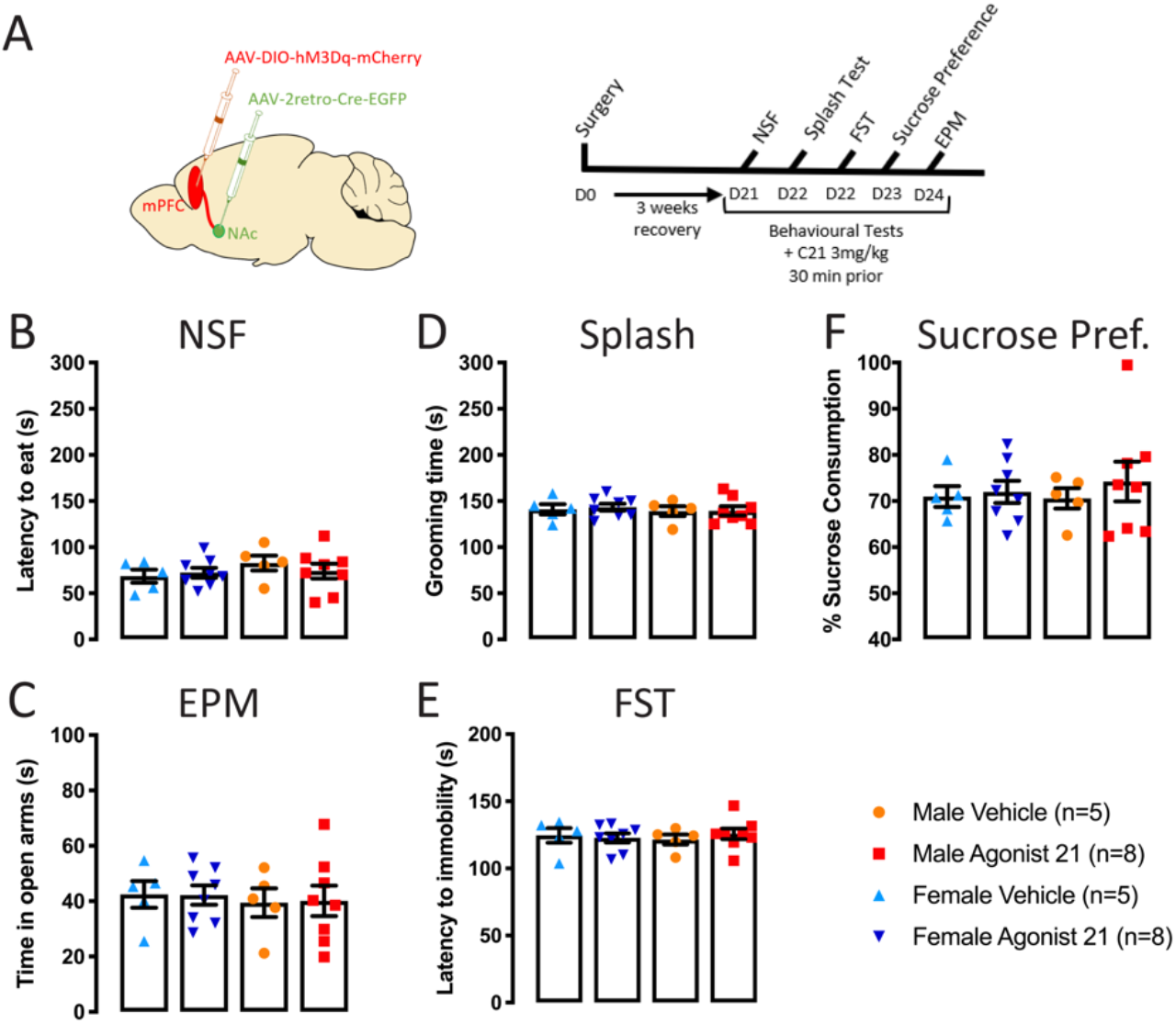
Acute activation of the NAc-projecting mPFC neurons is not sufficient to trigger depressive-like behaviours in males and females. **(A)** Schematic representation of the experimental timeline. **(B-F)** Chemogenetic activation of the NAc-projecting mPFC neurons in naive males and females is not inducing anxiety, behavioral despair or anhedonia as illustrated in the **(B)** NSF, **(C)** EPM, **(D)** splash test, **(E)** FST and **(F)** sucrose preference test. Significance was determined using two-way ANOVA with *post-hoc* Tukey multiple comparisons test. Symbols and bars represent the group average ± SEM.

## Tables

**Table 1.**
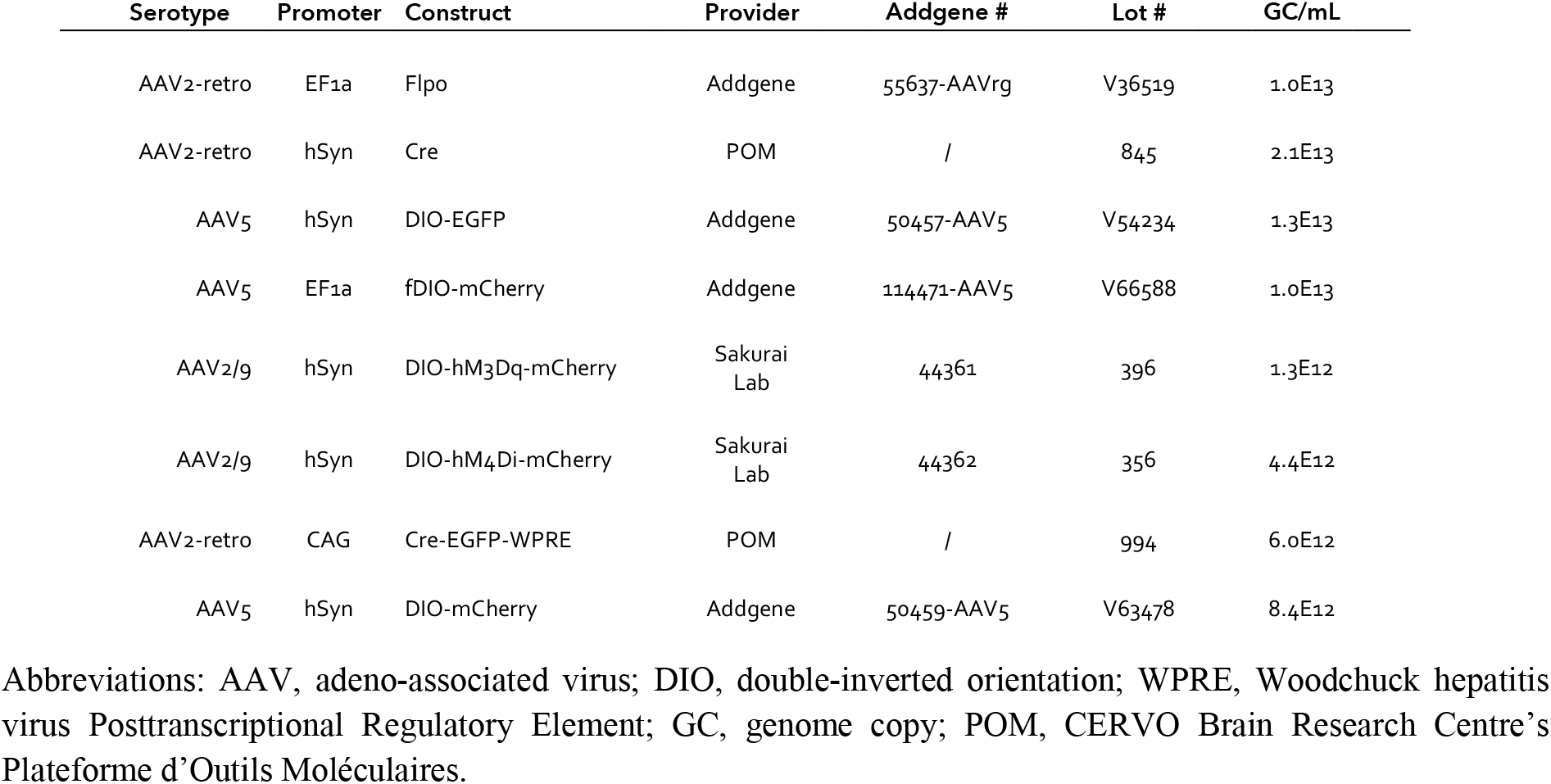
Adeno-associated viral vectors.

| Serotype | Promoter | Construct | Provider | Addgene # | Lot # | GC/mL |
| --- | --- | --- | --- | --- | --- | --- |
| AAV2-retro | EF1a | Flpo | Addgene | 55637-AAVrg | V36519 | 1.0E13 |
| AAV2-retro | hSyn | Cre | POM | / | 845 | 2.1E13 |
| AAV5 | hSyn | DIO-EGFP | Addgene | 50457-AAV5 | V54234 | 1.3E13 |
| AAV5 | EF1a | fDIO-mCherry | Addgene | 114471-AAV5 | V66588 | 1.0E13 |
| AAV2/9 | hSyn | DIO-hM3Dq-mCherry | Sakurai Lab | 44361 | 396 | 1.3E12 |
| AAV2/9 | hSyn | DIO-hM4Di-mCherry | Sakurai Lab | 44362 | 356 | 4.4E12 |
| AAV2-retro | CAG | Cre-EGFP-WPRE | POM | / | 994 | 6.0E12 |
| AAV5 | hSyn | DIO-mCherry | Addgene | 50459-AAV5 | V63478 | 8.4E12 |

Abbreviations: AAV, adeno-associated virus; DIO, double-inverted orientation; WPRE, Woodchuck hepatitis virus Posttranscriptional Regulatory Element; GC, genome copy; POM, CERVO Brain Research Centre's Plateforme d'Outils Moléculaires.

**Table 2.** Stereotaxic coordinates.

| Structure | Angle | AP | ML | DV | Volume |
| --- | --- | --- | --- | --- | --- |
| mPFC | 10° | 1.8 mm | ±0.75 mm | 2.3 mm | 0.5 µL |
| NAc | 10° | 1.55 mm | ±1.45mm | 4.4 mm | 0.3 µL |
| VTA | 10° | -3.2 mm | ±1.35 mm | 4.6 mm | 0.3 µL |

Abbreviations: mPFC, medial prefrontal cortex; NAc, nucleus accumbens; VTA, ventral tegmental area; AP, anteroposterior; ML, mediolateral; DV, dorsoventral.

**Table 3.**
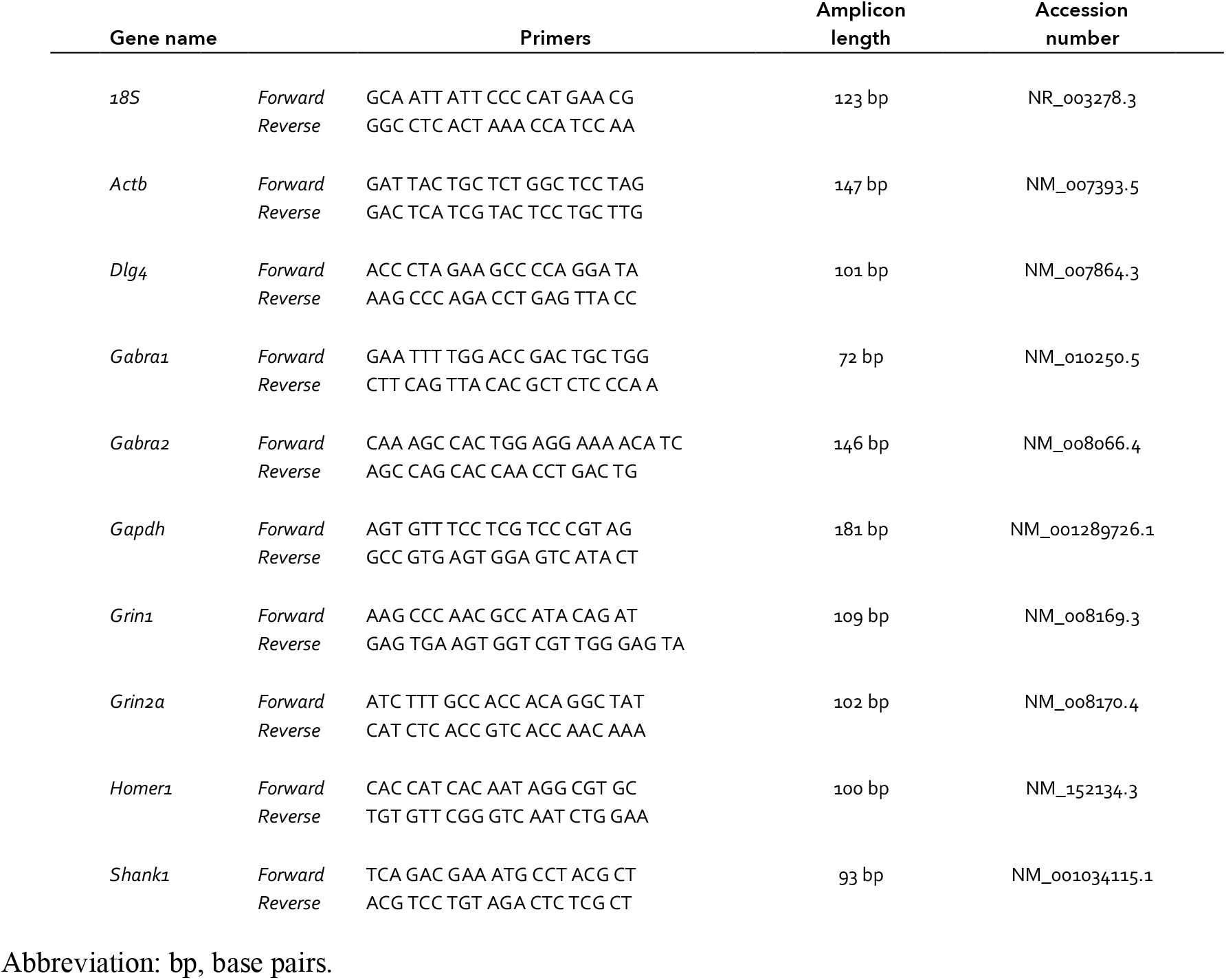
Primers for real-time quantitative PCR.

